# Integrated ERK, PKA, YAP/TAZ, and SHH Signaling Drives Cortical Lineage Diversification and Evolutionary Expansion

**DOI:** 10.1101/2025.04.09.647935

**Authors:** Zhuangzhi Zhang, Zhejun Xu, Tongye Fu, Jialin Li, Feihong Yang, Chuannan Yang, Wenhui Zheng, Zizhuo Sha, Yanjing Gao, Mengge Sun, Zhenmeiyu Li, Jing Ding, Xiaosu Li, Zhengang Yang

## Abstract

The human cerebral cortex, essential for intelligence, cognition, and language, evolved its uniqueness through mechanisms that drive neuronal expansion. Here, we demonstrate that ERK and PKA signaling cooperatively preserve the neurogenic capacity of cortical radial glia (RGs) by inhibiting gliogenic YAP/TAZ and SHH signaling. YAP/TAZ signaling drives cortical RGs toward ependymal fate, while SHH signaling facilitates the generation of cortical tripotential intermediate progenitor cells that produce cortical astrocytes, oligodendrocytes, and olfactory bulb interneurons. During evolution, cortical outer radial glia (oRGs) acquire dominant ERK/PKA signaling through a self-reinforcing feedback mechanism that suppresses both YAP and SHH pathways, thereby significantly enhancing oRG self-renewal capacity and extending the neurogenic period. These findings reveal that cortical neurogenesis, gliogenesis, and evolutionary expansion are coordinately regulated by an integrated ERK, PKA, YAP/TAZ, and SHH signaling, identifying a unifying principle of mammalian cortical development and evolution.

## 1. Introduction

Radial glial (RGs), spanning the entire thickness of the developing cortex, are primary neural stem cells ^1–8^ During mouse corticogenesis, three distinct types of RGs can be classified based on their cell lineage, which we designate as N-RGs, E-RGs, and T-RGs. The early RGs are neurogenic (N-RGs), generating intermediate progenitor cells (IPCs) exclusively for glutamatergic pyramidal neurons (PyNs), known as PyN-IPCs. ^1–8^ At the end of cortical neurogenesis, a subpopulation of N-RGs in the medial cortex transitions into E-RGs that produce ependymal cells along a medial-to-lateral gradient. ^9–12^ Meanwhile, another N-RG population transforms into T-RGs in a ventral-lateral-to-medial direction, giving rise to cortical tripotential intermediate progenitors (Tri-IPCs) that express ASCL1, EGFR, and OLIG1/2. ^4,13–18^

These Tri-IPCs sequentially generate cortical astrocyte lineage-restricted IPCs (APCs), oligodendrocyte lineage-restricted IPCs (OPCs), and IPCs for cortically derived olfactory bulb interneurons (OBIN-IPCs). These lineage-restricted IPCs then divide symmetrically to generate cortical astrocytes, oligodendrocytes, and OBINs, respectively. ^13–19^

During human corticogenesis, full-span radial glial cells (fRGs) persist from gestational week (GW) 8 through GW16. Around GW16, a large number of fRGs give rise to the truncated radial glia (tRGs) in the ventricular zone (VZ) and the outer radial glia (oRGs, also known as basal RGs) in the outer subventricular zone (outer SVZ, OSVZ). ^18,20–26^ Human cortical fRGs are neurogenic, mainly generating PyN-IPCs for deep layer PyNs. Human cortical oRGs, which are rare in mice, are also neurogenic, mainly generating PyN-IPCs for upper layer PyNs. ^4,15,17,18,20,25^

Similar to mouse cortical full-span RGs, human tRGs exhibit three distinct subtypes, referred to as N-tRGs, E-tRGs, and T-tRGs. After human N-tRGs are generated from fRGs at GW16, they start to express EGFR and ASCL1, and initially generating PyN-IPCs for upper layer PyNs (i.e., at GW16-GW17). ^14,18,27^ Subsequently, N-tRGs undergo a neurogenesis to gliogenesis transition. At approximately GW18, a subset of medial cortical N-tRGs transitions into E-tRGs, which subsequently generate the cortical ependymal cells. ^11,14,18,28^ In parallel, another population of lateral cortical N-tRGs transits into T-tRGs, giving rise to Tri-IPCs. ^14,15,18,25,27^ These Tri-IPCs give rise to human cortical APCs, OPCs, and cortically derived OBIN-IPCs. ^4,14,16,18^

Here, we elucidate the mechanisms governing lineage progression in cortical RGs. During cortical development, RGs adopt one of three fates, dictated by specific signaling pathways: (1) upregulation of ERK/PKA in N-RGs promotes sustained neurogenesis while suppressing gliogenesis; 2) activation of YAP/TAZ in E-RGs drives ependymal cell differentiation; and (3) enhanced SHH signaling in T-RGs produces cortical Tri-IPCs. During evolution, cortical oRGs developed dominant ERK/PKA signaling that inhibits YAP and SHH, boosting their self-renewal and prolonging neurogenesis. These findings demonstrate that cortical neurogenesis, gliogenesis, and evolutionary expansion constitute an integrated biological program coordinated by the conserved ERK, PKA, YAP, and SHH signaling. Our study reveals fundamental principles underlying cortical development and evolution.

## 2. Results

### 2.1 Molecular identity of human cortical oRGs, E-tRGs, and T-tRGs

Human cortical development exhibits a slower, more refined progression, offering a unique model to investigate the molecular mechanisms underlying neurogenesis, gliogenesis, and evolutionary expansion. To identify the molecular features of human cortical oRGs, E-tRGs, and T-tRGs, we analyzed scRNA-Seq data using the human GW22 cortical tissue. ^29^ The heatmap presents a comprehensive profile of 28 genes (**Figure** 1A, B), illustrating the molecular identity and potential cell lineage relationships among human cortical oRGs, E-tRGs, and T-tRGs (Figure 1C). Our analysis revealed that all pathways suppress SHH and YAP signaling (YAP/TAZ activity) in human cortical oRGs.

**Figure 1.**
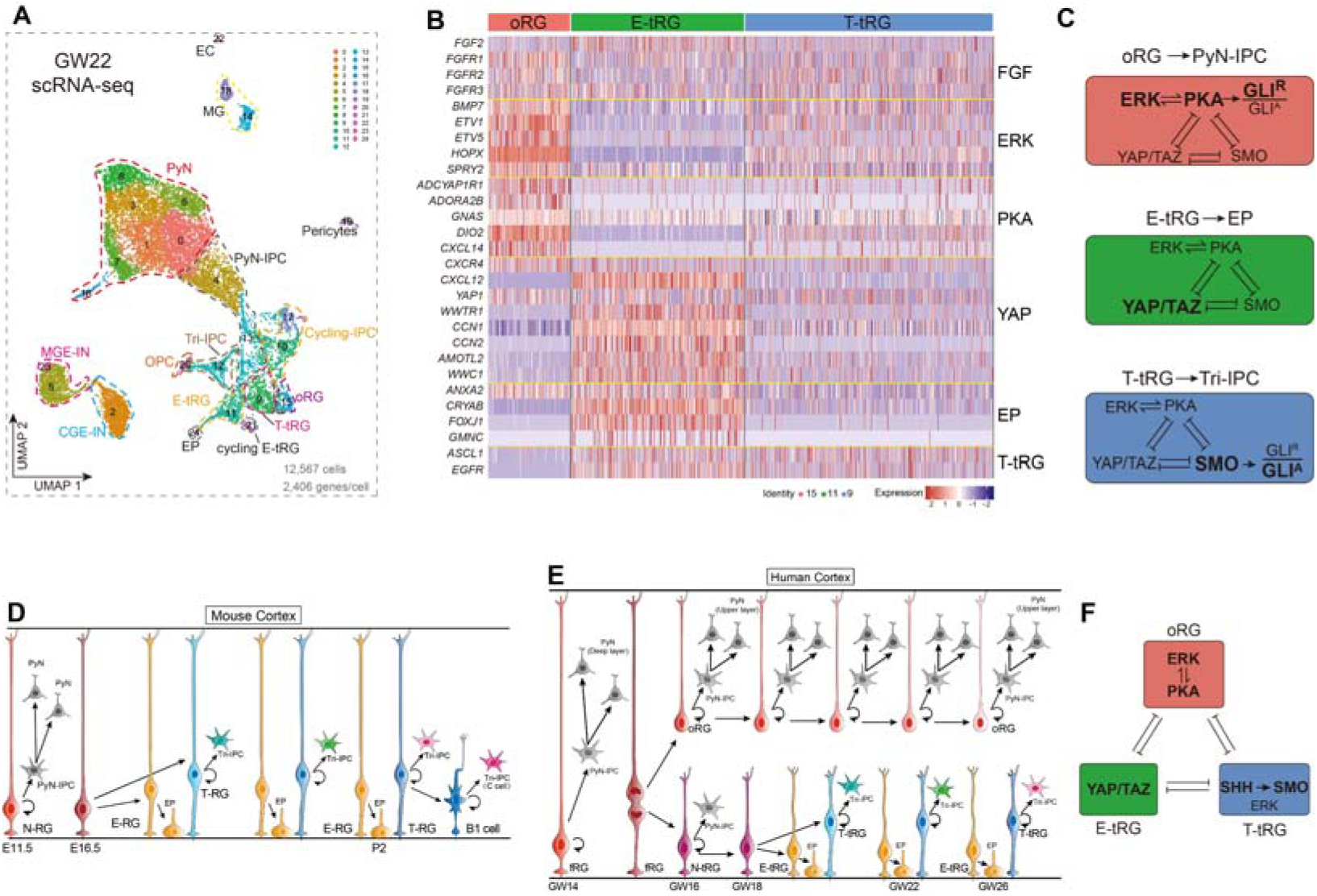
The molecular signatures of human cortical oRGs, E-tRGs, and T-tRGs. **A)** scRNA-Seq data derived from human cortical tissue at GW22. UMAP visualization with cells colored by cluster. **B)** Heatmap illustrating differentially expressed genes (DEGs) among human cortical oRGs, E-tRGs, and T-tRGs. EP, ependymal cells. **C)** oRGs exhibited elevated ERK and PKA signaling, while E-tRGs exhibited enhanced YAP signaling. A subset of T-tRGs expressed genes (such as *EGFR* and *ASCL1*) in response to elevated SHH-SMO signaling. **D, E)** Mouse and human cortical RG cell lineage progression. **F)** oRGs, E-tRGs, and T-tRGs preserve their distinct identities via coordinated evolutionarily conserved signaling pathways, including ERK, PKA, YAP/TAZ, and SHH signaling.

Extracellular signal-regulated kinase (ERK) signaling is a master regulator of cell behavior, governing nearly all cellular processes. ^30,31^ It can be activated through the binding of FGFs to FGFRs. ^17,30–34^ While *FGF2* and *FGFR1/2/3* are expressed at comparable levels in human cortical oRGs, E-tRGs, and T-tRGs, ERK signaling intensity varies, being strongest in oRGs, weakest in E-tRGs, and intermediate in T-tRGs (Figure 1B). This conclusion is based on the expression levels of ERK signaling response genes, including *BMP7, ETV1, ETV5, HOPX*, and *SPRY*2 (Figures 1B, and S1A, B, Supporting Information). ^17,25^ Moreover, we observed that cAMP-PKA signaling activity exhibits a pattern similar to that of ERK signaling across RGs. For example, G protein-coupled receptor genes *ADCYAP1R1* and *ADORA2B* are expressed much higher in oRGs than E-tRGs and T-tRGs (Figures 1B, and S1B, Supporting Information). ADCYAP1R1 binds to PyN-derived ADCYAP1, resulting in the activation of the coupled GNAS (Gsα) to stimulate adenylyl cyclase activity to increase cAMP-PKA signaling. ^35^ Likewise, adenosine binding to ADORA2B also activates cAMP-PKA signaling. ^36^ cAMP-PKA-CREB targets include *DIO2* ^37–39^ and *CXCL14*. ^40^ The DIO2 promoter in humans, rats, and mice contain a conserved canonical cAMP response element. ^37–39^ Accordingly, we observed that oRGs expressed higher levels of *DIO2* and *CXCL14* compared with E-tRGs and T-tRGs (Figures 1B, and S1B, Supporting Information), providing evidence that oRGs have high PKA activity. PKA-mediated phosphorylation of FGF2 increases its binding affinity up to 8-fold and potentiates ERK activity. ^41^ Moreover, ERK and PKA signaling can mutually reinforce each other. ^42–44^ PKA acts as a strong negative regulator of the SHH-SMO signaling, ^45–50^ and PKA is also widely recognized for its role in inhibiting YAP signaling. ^47,50–53^ Taken together, this analysis demonstrates that heightened ERK and PKA activity represents a defining feature of human cortical oRGs.

The Hippo pathway core kinases MST1/2 and LATS1/2 suppress cell growth by phosphorylating YAP/TAZ, inhibiting their nuclear translocation and transcriptional activity. When localized in the nucleus, YAP/TAZ bind TEAD transcription factors to promote cell proliferation and maintain homeostasis. ^47,50–53^ A prominent feature of human cortical E-tRGs is the robust activation of YAP signaling (YAP/TAZ activity), which is required for cortical ependymal cell formation. ^54,55^ YAP signaling signature genes are highly expressed in E-tRGs, including, *CCN1 (CYR61), CCN2 (CTGF), AMOTL2*, *WWC1/2*, and *WWTR1* (TAZ) (Figures 1B, and S1C, Supporting Information). Consequently, markers for early ependymal cells, including *CRYAB, FOXJ1*, *ANXA2,* and *GMNC*, are also expressed in E-tRGs (Figures 1B, and S1C, Supporting Information). Furthermore, *CXCL12/CXCR* signaling is upregulated in E-tRGs compared to oRGs and T-tRGs (Figures 1B, and S1C, Supporting Information), which represses cAMP-PKA signaling. ^50,56,57^ Indeed, PKA and ERK signaling are extremely low in E-tRGs (Figure 1B). This observation aligns with previous studies demonstrating a reciprocal regulatory relationship between these pathways: while PKA activity inhibits YAP signaling, ^47,50–53^ YAP signaling conversely suppresses both cAMP-PKA and ERK signaling. ^58,59^ Furthermore, YAP signaling inhibits primary cilium assembly, ^60^ which suppress SHH-SMO signaling. Collectively, this analysis provides evidence that YAP signaling, by repression of PKA, ERK, and SHH signaling, is a key feature of E-tRGs.

At GW22, T-tRGs exhibit low to moderate levels of ERK, PKA, YAP, and SHH signaling (Figure 1B, and S1A-D, Supporting Information), indicating the convergence of multiple signaling pathways. Indeed, *EGFR* and *ASCL1*, which are activated by ERK, SHH and/or YAP signaling, are expressed by a subpopulation of T-tRGs. ^15,16,18,25,27,61^ These T-tRGs give rise to Tri-IPCs that subsequently generate APCs, OPCs and OBIN-IPCs. ^4,14,15,18,25^ However, at GW22, we identified APCs and OPCs, but not OBIN-IPCs, derived from Tri-IPCs (Figure S2A-F, Supporting Information). With increased SHH signaling in the cortical VZ at GW23 and GW26, there was increased expression of *GLI1* and *HHIP* (SHH pathway markers) in T-tRGs and their progenies. Furthermore, genes of OBIN-IPCs, including *GSX2, DLX2, GAD2*, and *SP9*, became evident within the Tri-IPC cluster (Figure S3A-G, Supporting Information). SHH-SMO signaling represses PKA signaling. ^45,46,49,50^ The SHH-SMO signaling may additionally suppress ERK signaling through the activation of PP2A-mediated mechanisms. ^62,63^ Studies also suggested that the functional interplay between YAP and SHH signaling governs the specification of different cell types. ^64^ As observed in the cortex, YAP signaling drives the cortical E-tRG-derived multiciliated ependymal cell lineage, ^54,55^ whereas SHH-SMO signaling promotes the primary-ciliated T-tRG-derived Tri-IPC cell lineage. ^13,15,17,25^

In summary, during the later stages of human cortical development, PyN-IPC-generating oRGs, ependymocyte-IPC-generating E-tRGs, and Tri-IPC-generating T-tRGs develop distinct molecular signatures, driven by different conserved signaling pathways that mutually inhibit one another (Figure 1C-F). These findings reveal that ERK and PKA signaling synergistically sustain prolonged self-renewal and neurogenic capacity in human oRGs, a critical driver of cortical neurogenesis and expansion, by coordinately suppressing the gliogenic YAP/TAZ and SHH pathways. To experimentally validate these mechanistic insights, we employed complementary mouse genetic approaches.

### 2.2 YAP signaling controls cortical E-RG identity

Increased nuclear translocation of YAP/TAZ proteins in mouse cortical RGs from E12.5 to E16.5, enhances YAP/TAZ activity, which correlates with advancing stages of cortical development. ^54,55^ To investigate the role of YAP signaling in cortical RGs, we crossed mice harboring a *WWC*-derived floxed minigene (referred to as *SuperHippo*) with *hGFAP-Cre* mice. *SuperHippo* specifically inhibits YAP signaling, mimicking *Yap1/Wwtr1* double knockout mice (**Figure** 2A). ^65^ To label E15.5 progenitors, we injected FlashTag into the mouse lateral ventricle. At E16.5, we isolated FlashTag-labeled cortical cells for scRNA-Seq analysis (Figure 2B). Blocking YAP signaling reduced expression of *Cyr61, Ctgf, Amotl2*, and *Wwc2* in cortical RGs, and eliminated expression of *Cryab* and *Foxj1* (early ependymal cell markers) (Figure 2C, D). On the other hand, ERK signaling response genes (e.g. *Etv1, Etv5, Spred1/*2, *Spry2, Bmp7, Id1/3*, and *Hopx*) were upregulated (Figure 2D). *Gnas*, a signature PKA signaling gene, was also increased, suggesting that a subpopulation of cortical RGs in *hGFAP-Cre; SuperHippo* mice exhibit molecular features of human oRGs, based on their higher ERK and PKA signaling (Figure 2D).

**Figure 2.**
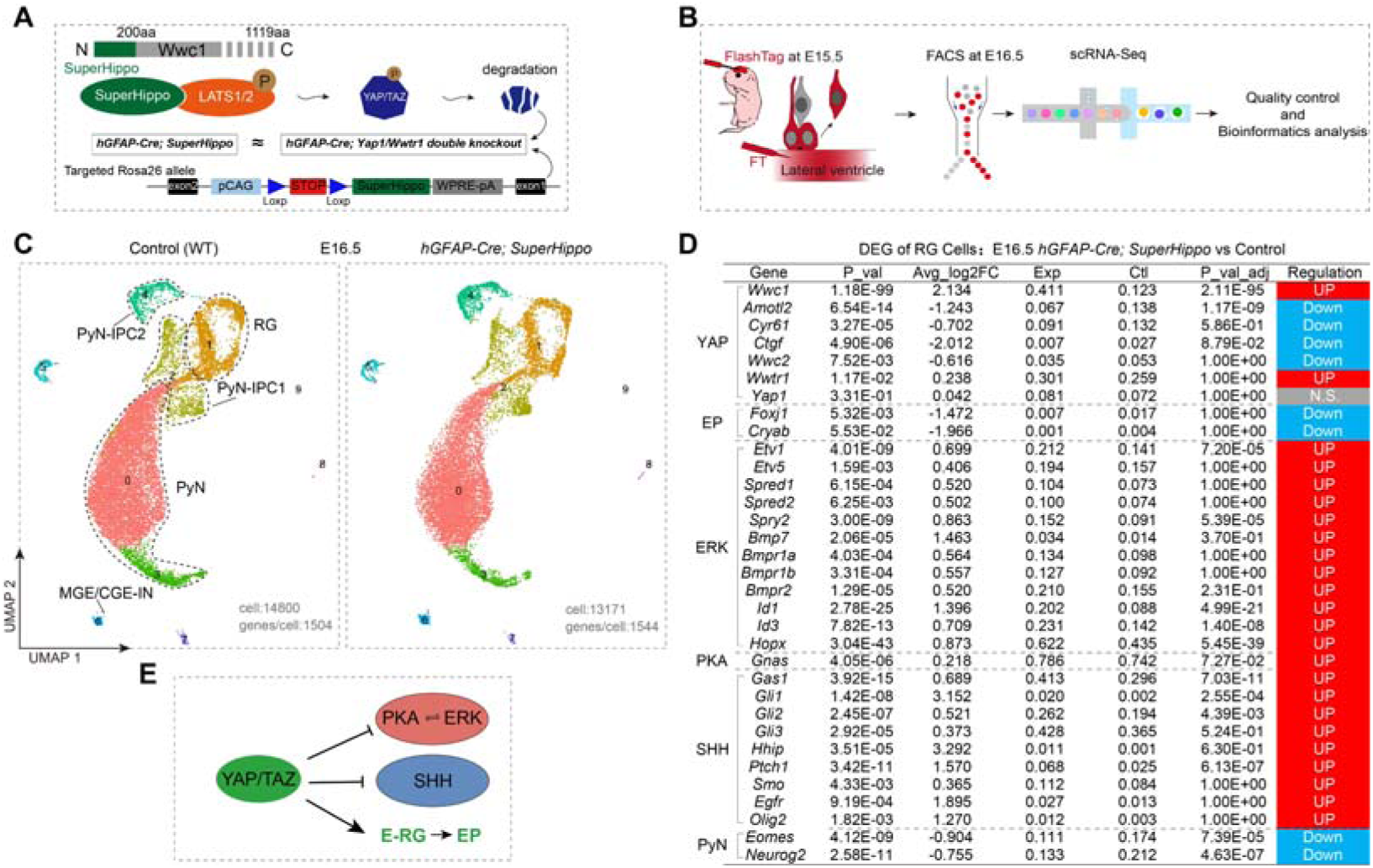
YAP signaling promotes the cortical E-RG identity. **A)** A *SuperHippo* minigene has been engineered to robustly inhibit YAP signaling. **B)** Strategy for labeling (lateral ventricle injection of FlashTag-CellTrace Yellow at E15.5) and enriching cortical progenitor cells from two littermate controls (wild type, WT) brains and two *hGFAP-Cre; SuperHippo* brains at E16.5 and scRNA-Seq analysis. **C)** Annotation of cortical cell clusters. **D)** scRNA-Seq analysis revealed DEGs in E16.5 cortical RGs (cluster 1 in A) of *hGFAP-Cre; SuperHippo* mice relatively to WT controls. **E)** YAP signaling governs cortical E-RG identity maintenance and ependymal differentiation by coordinately inhibiting PKA/ERK activity and suppressing SHH singling.

The *hGFAP-Cre; SuperHippo* mutant had increased SHH-SMO signaling based in upregulation of *Gas1, Gli1, Gli2, Gli3, Hhip, Ptch1*, *Smo,* and *EGFR* (Figure 2D), suggesting that a subpopulation of cortical RGs in *hGFAP-Cre; SuperHippo* mice had molecular features of T-RGs. ERK signaling enhances self-renewal of cortical RGs,^17,66–68^ while SHH-SMO signaling promotes the generation of Tri-IPCs from cortical RGs. ^13,15^ Correspondingly, expression of PyN-IPC marker genes *Eomes* (*Tbr2*) and *Neurog2* was downregulated in the mutant cortical RGs (Figure 2D).

By P2, E-RGs and their ependymal cell progeny were absent in *hGFAP-Cre; SuperHippo* mice (**Figure**s 3A, B, and S4A, B, Supporting Information), consistent with prior reports demonstrating the essential role of YAP signaling in ependymal cell development. ^54,55^ Immunohistochemical analysis demonstrated an increase in Tri-IPCs (OLIG2- and GSX2-expressing cells) and the loss of ependymal cells (Figure 3C-E). Although cortical E-RGs and ependymal cells are lost in *hGFAP-Cre; SuperHippo* mice, cortical T-RGs were preserved but still showed downregulation of YAP-signaling genes, and the expression of *Cryab, Foxj1*, and *Ogn* (ependymal markers), was abolished (Figures 3A-C, and S4B, Supporting Information). In contrast, ERK, PKA, and SHH signaling were upregulated (Figures 3B, and S4, Supporting Information). All of these findings were further confirmed via scRNA-Seq and immunohistochemical analysis of *Emx1-Cre; SuperHippo* mice (Figure S5A-E, Supporting Information).

**Figure 3.**
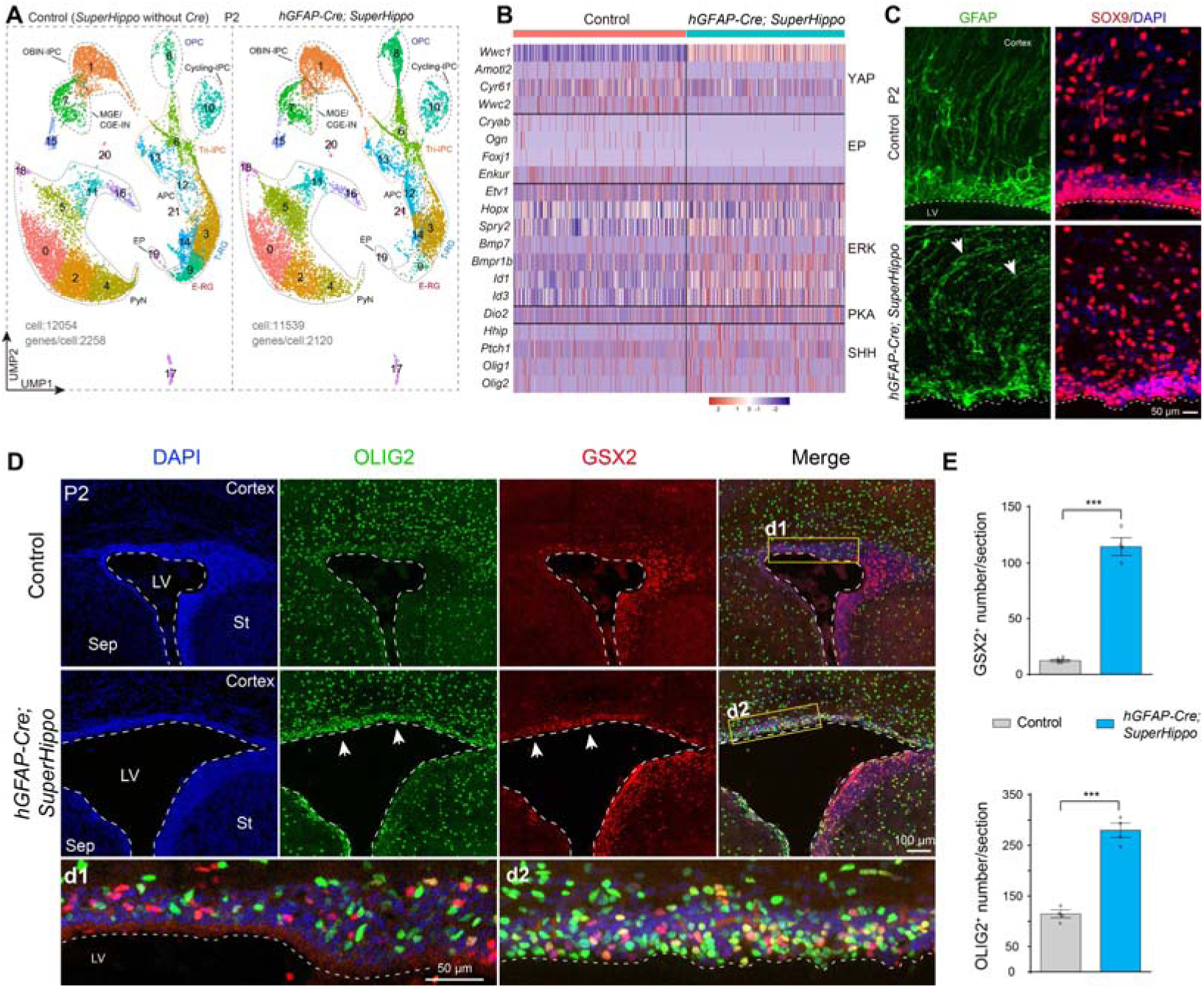
YAP signaling serves as a master regulator of cortical ependymal cell production from RGs. **A)** Annotation of cortical cell clusters in P2 control (*SuperHippo* heterozygotes without *hGFAP-Cre*) and *hGFAP-Cre; SuperHippo* mice. **B)** Heatmap illustrating DEGs in cortical T-RGs (cluster 3 in **A**). **C)** Immunostaining reveals disruption of the cortical ventricular surface and the loss of apical SOX9- and GFAP-expressing immature ependymal cells in the P2 *hGFAP-Cre; SuperHippo* cortex. LV, lateral ventricle. **D, E)** Immunostaining reveals disruption of the cortical ventricular surface and increase in OLIG2- and GSX2-expressing progenitors in the cortical VZ/SVZ of *hGFAP-Cre; SuperHippo* mice at P2 (*n* = 4 per group; mean ± SEM; \*\*\**P* <0.001; unpaired two-tailed Student’s *t*-test).

Finally, we reanalyzed bulk RNA-Seq and ChIP-Seq data from mice expressing the YAP1-MAMLD1 (YAP-M) fusion protein, which is implicated in supratentorial ependymoma. ^61^ The gene expression profiles demonstrated distinct patterns between cells from the SVZ of wild-type mice (n=3) and tumor cells derived from ∼P20 mice transfected with YAP-M at E14.5 (n=4) (Figure S6A, Supporting Information). The YAP-M fusion oncogene significantly upregulated expression of YAP signaling components and early ependymal cell markers (Figure S6A, Supporting Information). Conversely, YAP-M oncogene exhibited a strong suppressive effect on expression key signaling pathway genes, including ERK, PKA, and SHH cascades, in addition to genes characteristic of cortical progenitor populations such as Tri-IPCs, APCs, OPCs, and OBIN-IPCs (Figure S6A, Supporting Information). This is consistent with observations that human YAP-M-driven tumor cells lack classical glial progenitor markers. ^61,69^ YAP1 and WWTR1 are transcriptional co-activators. Reanalysis of ChIP-Seq and Cut&Run data for YAP1 and H3K27ac in both YAP-M-induced tumor cells and YAP1-expressing mouse neural stem cells identified multiple YAP-M target genes, characterized by distinct YAP-M binding sites, mainly in the promoter region. ^70^ These direct target genes included *Foxj1, Cryab, Anxa2, S100a11*, and *Ogn*, among others (Figure S6B, C, Supporting Information). Taken together, these loss- and gain-of-function analyses demonstrate that YAP signaling promotes cortical E-RG identity during cortical development and suppresses ERK, PKA, and SHH signaling (Figure 2E).

### 2.3 SHH-SMO drives the establishment of cortical T-RG identity

In all studied animals, a primary function of active SMO is to counteract the inhibitory effects of PKA on GLI transcription factors. ^45^ To investigate the *Smo* function in cortical RGs, we reanalyzed our previous scRNA-Seq data of cortical cells in *hGFAP-Cre; Smo^F/F^* mice E18.0. ^25^ In the absence of *Smo*, cortical RG expression of *Gli1, Hhip, Ptch1, Egfr*, and *Olig2* was nearly undetectable (**Figure** 4A-C). In contrast, signature genes associated with ERK signaling (*Hopx, Bmp7, Etv5*, etc.), PKA signaling (*Adcyap1r1, Adora2b, Gnas, Dio2*, and *Cxcl14*), and early ependymal cell markers (*Anxa2, Aqp4, Cryab, Foxj1*, and *S100a6*) exhibited significant upregulation (Figure 4C). Notably, the observed upregulation of YAP signaling-responsive genes was not pronounced (Figure 4C). It is likely that this modest change in expression at E18 is due to the potent inhibitory effect of PKA signaling on YAP activity. Additionally, the intensity of ventral-derived SHH-SMO signaling exhibits an inherently low level prenatally, which increases significantly after birth in mice. ^71^ To further validate these findings, we conducted additional scRNA-Seq analysis of cortical cells isolated from *hGFAP-Cre; Smo^F/F^* mice and their *Smo^F/^*^F^ littermate controls at P2. Notably, both E-RGs and T-RGs exhibited upregulation of ERK, PKA, and YAP signaling pathways (Figure S7A-C, Supporting Information). Taken together, these findings indicate that SHH-SMO signaling promotes and maintains mouse cortical T-RG identity and facilitates their differentiation into Tri-IPCs by suppressing ERK, PKA, and YAP signaling (Figure 4F).

**Figure 4.**
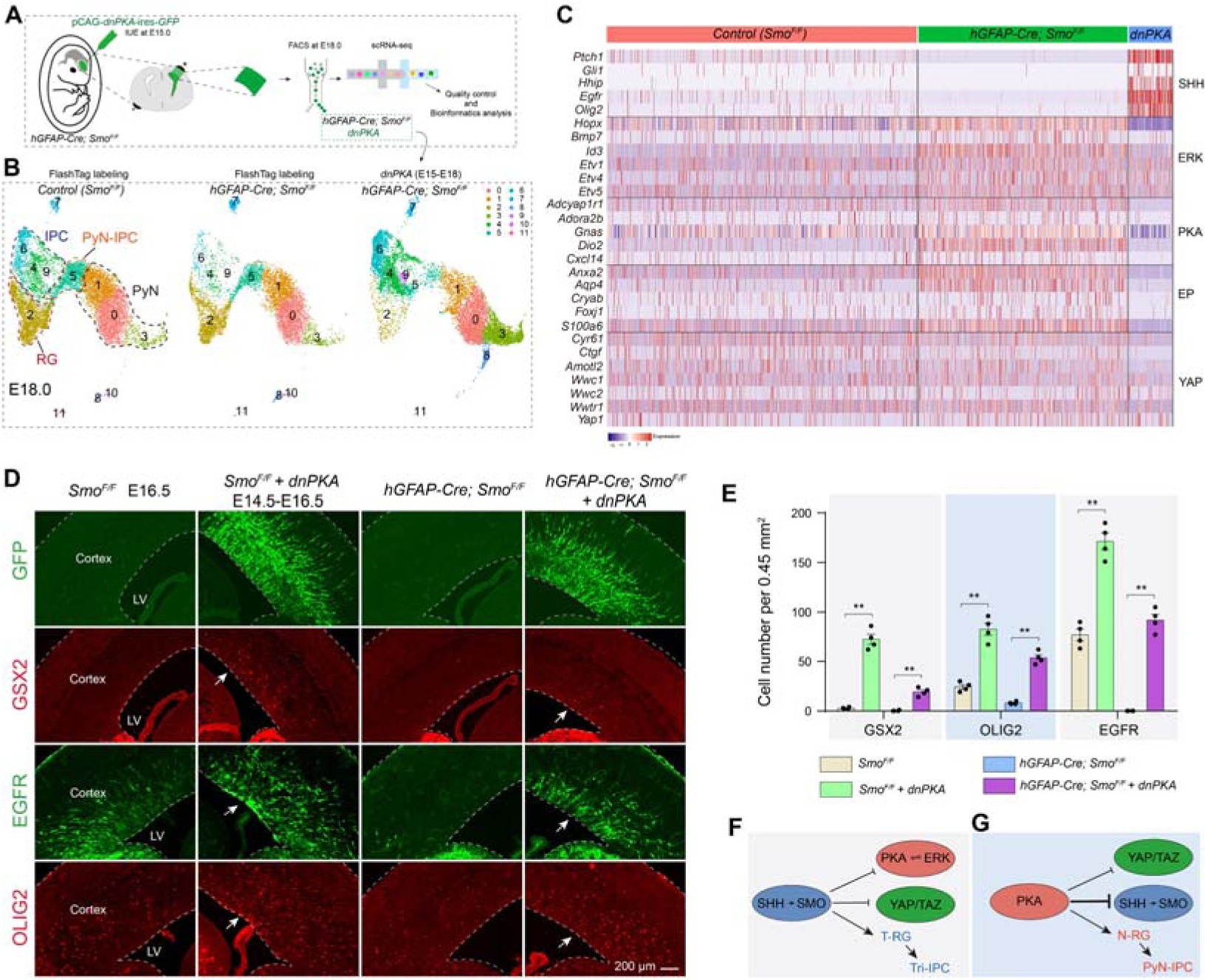
The functional interplay between SHH and PKA signaling in the cortical RGs. **A)** Scheme for overexpressing *pCAG-dnPKA-GFP* plasmids into the cortical VZ of *Smo^F/F^* control and *hGFAP-Cre; Smo^F/F^* mouse embryos at E15 by IUE, and scRNA-Seq of GFP-expressing cells at E18.0. **B)** Annotation of cortical cell clusters. **C)** Heatmap illustrating gene expression in cortical RGs with IUE of *dnPKA*. **D, E)** Immunostaining reveals a marked expansion of GSX2-, EGFR-, and OLIG2-positive progenitors in the cortical VZ/SVZ following IUE of *dnPKA* (PKA signaling completely ablation) (*n* = 4 per group; mean ± SEM; \*\**P* <0.01; one-way ANOVA and Tukey’s post-hoc test). **F)** SHH-SMO signaling promotes the identity of T-RGs. **G)** PKA signaling maintains cortical N-RG identity by strongly inhibiting SHH signaling and moderately suppressing YAP signaling.

### 2.4 PKA protects cortical N-RG identity

The reciprocal regulatory relationship between SHH signaling and PKA activity is well characterized: elevated PKA activity suppresses SHH signaling, while reduced PKA activity potentiates SHH pathway activation. ^45,46,49^ To further investigate the functional role of PKA in cortical RGs under conditions of *Smo* deficiency, we next performed a series of genetic analyses. At E15, we conducted IUE to ectopically express a dominant-negative form of PKA (d*nPKA-GFP*) in the mouse cortex. By E18, cells expressing *dnPKA-GFP* were isolated and analyzed by scRNA-Seq. Comparative analysis with FlashTag labeled cells from control (*Smo^F/F^*) and *hGFAP-Cre; Smo^F/F^* E18 mice without *dnPKA*-IUE revealed the following findings in the cortical RGs of *hGFAP-Cre; Smo^F/F^* mice expressing *dnPKA* (Figures 4A-C, and S8A, B, Supporting Information): (1) complete ablation of PKA signaling; (2) significant upregulation of SHH pathway genes; (3) marked downregulation of ERK signaling; (4) suppression of YAP signaling; (5) blockade of early ependymal cell marker genes. Thus, the complete loss of PKA in cortical RGs directly leads to a significant increase in SHH signaling, even without *Smo* function, which subsequently represses both ERK and YAP signaling. This result closely mirrors the phenotypes observed in our prior studies involving the overexpression of *Shh-N*, *Gli1*, *Gli2 activator*, or *SmoM2* in cortical RGs. ^13,15,17,25^ Our results further solidify the role of PKA as the principal negative regulator of SHH signaling, ^45,46,49^ in addition to its role as a suppressor of YAP signaling. ^50,51^ Immunostaining analysis of cortical sections 48-hours after *dnPKA* IUE revealed upregulated expression of EGFR, OLIG2, and GSX2 in the cortex of control and *hGFAP-Cre; Smo^F/F^*mice (Figure 4D, E). This highlights the pivotal role of PKA signaling in N-RGs (neurogenic RGs) in protecting PyN genesis while simultaneously suppressing gliogenesis, including the production of ependymal cells, astrocytes, oligodendrocytes, and OBINs (Figure 4G).

Therefore, genetic perturbations, including loss of SHH function (*hGFAP-Cre; Smo^F/F^*), gain of SHH function, and loss of PKA function (both via *dnPKA* overexpression), further support our model. This model posits that a tripartite signaling network, centered on cross-repressive interactions among the ERK/PKA, YAP, and SHH pathways, regulates mammalian cortical neurogenesis, ependymal gliogenesis, and macroglial (Tri-IPC) genesis (Figure 4F, G).

### 2.5 ERK safeguards both N-RG and T-RG identity

ERK signaling regulates multiple steps in cortical development. ^15,30–34^ To examine the function of ERK signaling, we used both *Emx1-Cre* and *hGFAP-Cre* lines to conditionally delete *Map2k1* and *Map2k2* (core downstream components of ERK; *Map2k1/2-dcko*) at progressively later stages. ^15,17^ scRNA-Seq analysis of E15.0 cortices revealed that SHH co-receptor *Gas1* expression was downregulated (**Figure** 5A, B), and expression of ERK signaling downstream target genes was completely lost in N-RGs (Figure 5B). This results in a significant decrease in the population of cortical N-RGs, concurrent with elevated *Eomes* expression (Figure 5B). These cellular alterations were associated with pronounced microcephaly, as evidenced by a substantial reduction in overall brain size (Figure S9A-C, Supporting Information), again demonstrating that ERK signaling in cortical N-RGs promotes self-renewal through enhanced cell proliferation while simultaneously inhibiting the generation of PyN-IPCs. ^17,66–68^ The upregulation of *Nr2f1* and *Fgfr3* expression in N-RGs confirmed that ERK signaling is essential for promoting the rostral cortical identity (Figure 5B). ^17,32^ Most importantly, we observed a subpopulation of cortical N-RGs exhibited *Adcyap1r1* expression, which was markedly downregulated in *Emx1-Cre; Map2k1/2-dcko* mice (Figure 5B). Conversely, there was upregulation of *Pde4d* expression in N-RGs (Figure 5B). ERK-mediated phosphorylation of PDE4D3 at Ser579 inhibits PDE4 enzymatic activity, thereby potentiating cAMP-PKA signaling pathways. ^44^ Consequently, ERK deficiency initiates a signaling cascade characterized by attenuated PKA activity, leading to YAP signaling activation (upregulation of *Ctgf*, *Amotl2*, and *Wwtr1*) and elevated expression of early ependymal cell marker genes (*Anxa2*, *Cryab*, and *Foxj1*) in cortical RGs of *Emx1-Cre; Map2k1/2-dcko* mice at E15.0 (Figure 5B).

**Figure 5.**
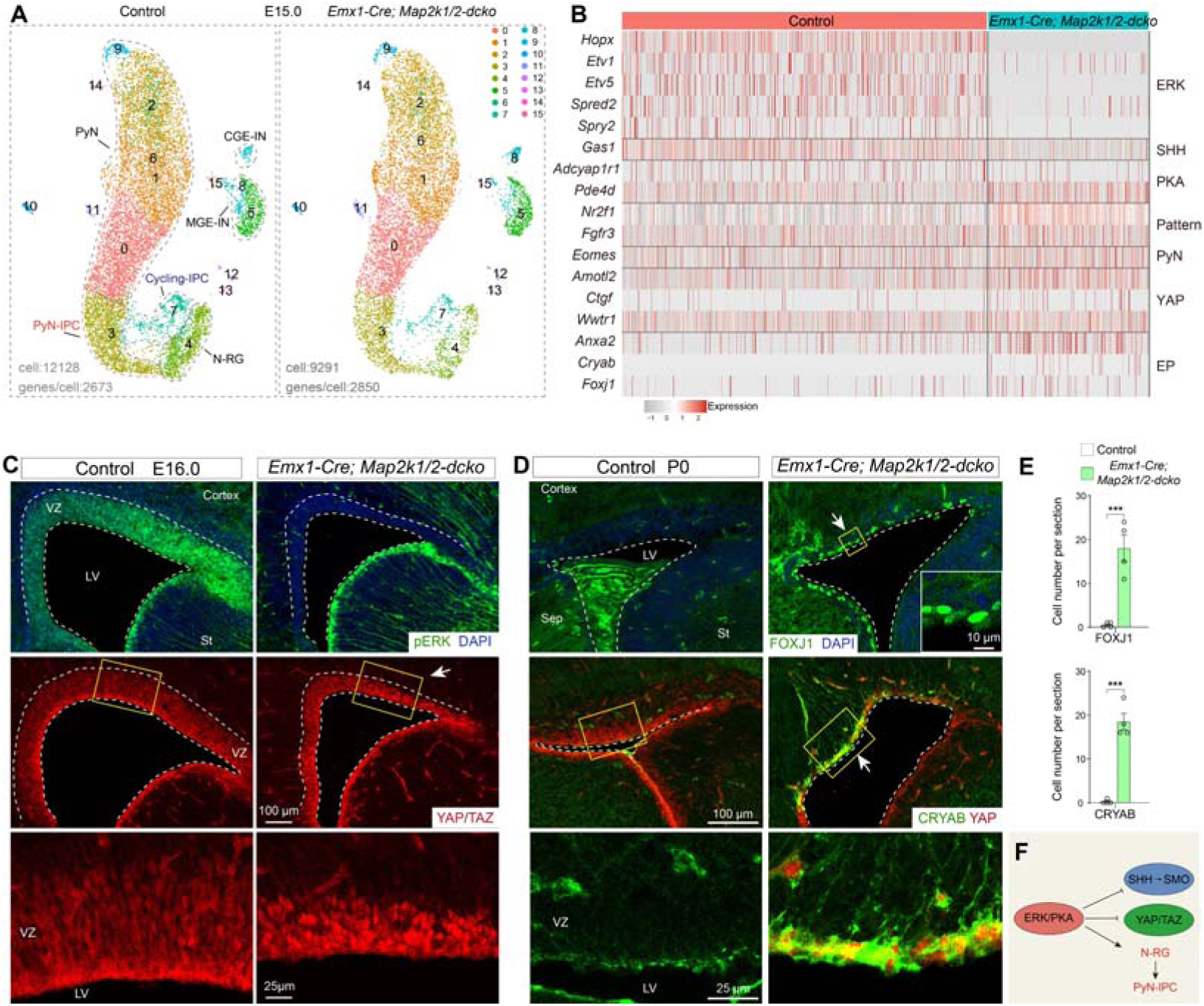
ERK signaling enhances PKA signaling and suppresses YAP signaling. **A)** UMAP showing annotated cell clusters based on scRNA-Seq of E15.0 control and *Emx1-cre; Map2k1/2-dcko* cortical cells. N-RG: neurogenic RG. **B)** Heatmap depicting DEGs across selected ERK, SHH, PKA, and YAP signaling response genes, along with cortical regional patterning markers and E-RG genes within cortical N-RGs. **C)** pERK (phosphorylated ERK) staining is lost, and nuclear localization of YAP/TAZ proteins is increased specifically in cortical RGs of *Emx1-Cre; Map2k1/2-dcko* mice at E16.0. **D, E)** FOXJ1- and CRYAB-expressing cells are observed in the VZ of *Emx1-Cre; Map2k1/2-dcko* mice, whereas their expression is undetectable in controls at P0 (*n* = 4 per group; mean ± SEM; \*\*\**P* <0.001; unpaired two-tailed Student’s *t*-test). Sep, septum; St, striatum. **F)** ERK/PKA signaling maintains the neurogenic program of cortical N-RGs by repressing gliogenic YAP and SHH signaling.

To further investigate the conserved functions of ERK signaling, we conducted a comprehensive reanalysis of our scRNA-Seq datasets from both *Emx1-Cre; Map2k1/2-dcko* and *hGFAP-Cre; Map2k1/2-dcko* mice at P2. ^15^ While *Emx1-Cre; Map2k1/2-dcko* mice displayed a substantial depletion of cortical RGs (**Figure** 6A, B), both mutants shared RGs defects, including: (1) loss of ERK signaling; (2) reduction of PKA signaling; (3) attenuation of SHH-SMO signaling; (4) failure to generate Tri-IPCs; (5) activation of YAP signaling; (6) upregulation of early ependymal cell marker gene expression; and (7) increased production of ependymal cell populations (Figures 6A, B, and S10A-C, Supporting Information). Our immunohistochemistry analyses yielded consistent results: a significant increase in YAP/TAZ nuclear localization within the cortical VZ at E16.0 (Figure 5C), followed by elevated numbers of CRYAB- and FOXJ1-expressing cells at P0 and P2 in *Emx1-Cre; Map2k1/2-dcko* mice compared to control littermates (Figures 5D, E, and 6C, D). We next employed a gain-of-function approach using *Emx1-Cre; Mek1DD* mice, which specifically enhances ERK signaling in cortical RGs starting from E10.5. This resulted in a marked expansion of the cortical RG population while simultaneously inhibiting neuronal differentiation, ultimately leading to pronounced macrocephaly (Figure S9D-H, Supporting Information).

**Figure 6.**
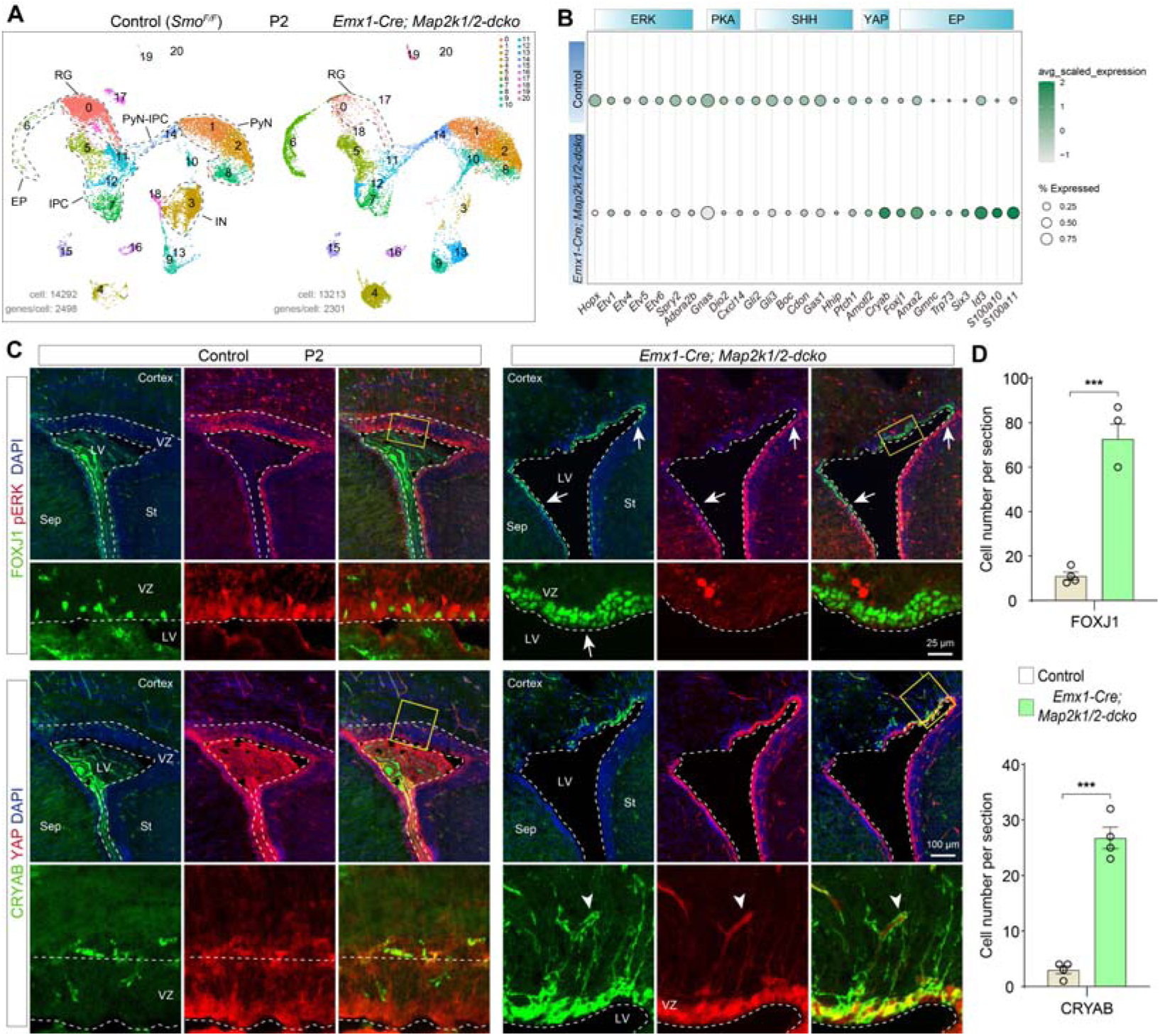
Loss of ERK signaling leads to downregulated PKA and SHH signaling, and increased YAP signaling. **A)** UMAP showing annotated cell clusters based on scRNA-Seq analysis of P2 control and *Emx1-cre; Map2k1/2-dcko* cortical cells. **B)** Bubble plot displaying DEGs among selected ERK, PKA, SHH, and YAP signaling response genes, as well as early cortical ependymal cell markers, in cortical RGs (cluster 0 in **A**) of *Emx1-Cre; Map2k1/2-dcko* mice compared controls at P2. **C, D)** FOXJ1- and CRYAB-expressing cells are significantly increased in the cortical VZ of *Emx1-Cre; Map2k1/2-dcko* mice, compared to the few FOXJ1- and CRYAB-expressing cells observed in control mice at P2 (*n* = 4 per group; mean ± SEM; \*\*\**P* <0.001; unpaired two-tailed Student’s *t*-test). Note that, in blood vessels, endothelial cells and vascular smooth muscle cells also expressed CRYAB and YAP/TAZ (arrowheads).

In *Emx1-Cre; Map2k1/2-dcko* mice, the degradation of pERK downregulates ERK and PKA signaling. While basal PKA activity potently inhibits ventrally-derived SHH-SMO signaling, it is insufficient to counteract local YAP activation, which occurs earlier than SHH-SMO signaling in cortical RGs. Indeed, low-level YAP/TAZ activity, activated by non-canonical WNT signaling along the cortical midline, was detected as early as E10.5 in cortical RGs. ^54,72–74^ This early patterned expression establishes a foundation for the medial-high to lateral-low gradient of YAP signaling observed in later stages. This temporal signaling hierarchy (elevated YAP signaling) directs cortical N-RGs to initially produce E-RGs, facilitating ependymal cell formation while inhibiting SHH signaling. Consequently, in the *Map2k1/2-dcko,* there is a block of N-RGs to differentiate into T-RGs and their Tri-IPC progeny. ^15^ However, experimental enhancement of SHH signaling in cortical RGs of *Emx1-Cre; Map2k1/2-dcko* mice, which exhibit repressed YAP signaling, rescued Tri-IPC generation. ^15^ In summary, during cortical neurogenesis, ERK/PKA signaling cooperate to suppress gliogenic YAP/TAZ and SHH signaling, thereby maintaining RG neurogenic potential (Figure 5F). During cortical gliogenesis, however, ERK/PKA signaling in the lateral cortex acts to suppress YAP/TAZ activity, thereby safeguarding the T-RG identity and ensuring the generation of Tri-IPCs. ^15^ This protective effect arises because SHH-SMO signaling more readily inhibits PKA than it does YAP signaling.^45–50^

## 3. Discussion

During the early regional patterning stage of corticogenesis, the development is predominantly orchestrated by WNT, FGF, and BMP signaling. ^32,75,76^ This signaling network promotes RGs to generate PyNs through a precisely regulated inside-out developmental sequence. ^1–5,77^ Next, the neurogenic period of cortical RGs is regulated by a combination of intrinsic molecular mechanisms and extrinsic environmental cues (Figure 7). In mice, as YAP and SHH signaling pathways intensify in cortical RGs, they antagonize ERK and PKA signaling, reaching a critical threshold around E16.5. At this stage, a subset of medial N-RGs transitions into E-RGs, which subsequently differentiate into ependymal cells. ^9–12^ Concurrently, N-RGs in the lateral cortex differentiate into T-RGs, initiating the production of Tri-IPCs (Figure 7A). ^15,16,18^

**Figure 7.**
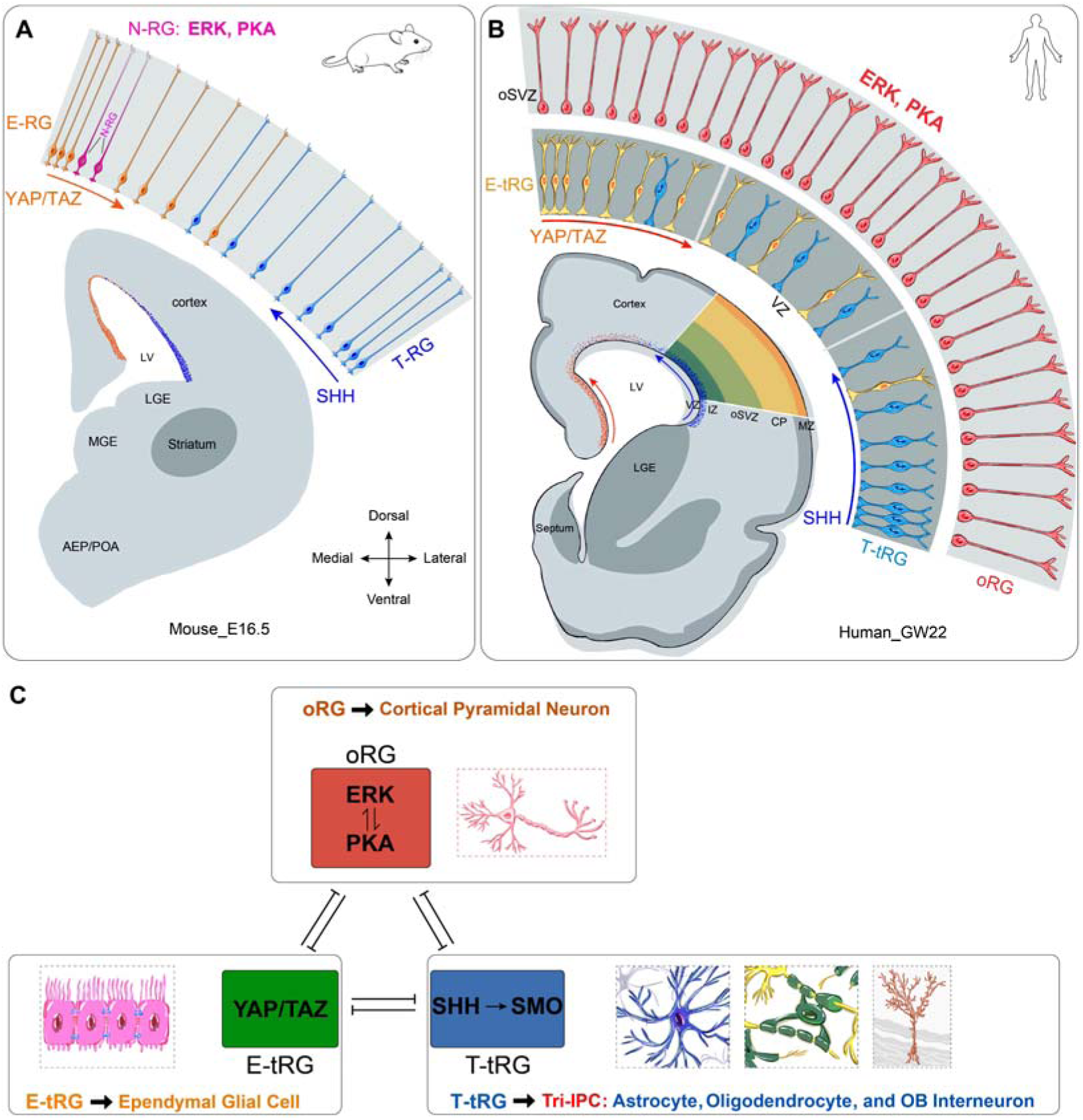
The integrated ERK, PKA, YAP/TAZ, and SHH signaling drives mammalian cortical expansion and lineage diversification. **A)** Cortical RG subtypes in mouse at E16.5. **B)** At GW22, human cortical RGs primarily consist of: oRGs in the OSVZ, generating upper-layer PyNs; E-tRGs in the VZ, producing cortical ependymal glial cells; T-tRGs in the VZ, giving rise to Tri-IPCs, which sequentially differentiate into astrocytes, oligodendrocytes, and cortically derived olfactory bulb interneurons. **C)** During corticogenesis, cortical RGs adopt one of three distinct developmental trajectories, each regulated by specific signaling pathways: ERK/PKA upregulation in RGs results in sustained neurogenesis with concomitant suppression of gliogenesis; YAP/TAZ upregulation in RGs results in generating ependymal cells; enhanced SHH signaling in RGs results in generating Tri-IPCs. Note that during cortical evolution, ERK/PKA signaling becomes the dominant pathway in cortical oRGs via a self-reinforcing feedback loop that actively suppresses both YAP and SHH signaling. This evolutionary adaptation confers two critical advantages: an enhanced self-renewal capacity in oRGs and a prolonged neurogenic period in RGs. More importantly, FGF-ERK signaling exhibits a rostral-high to caudal-low gradient in the cortex, which promotes the expansion of the prefrontal cortex. Collectively, these mechanisms drive the dramatic expansion of neuronal output, neocortical volume, and the prefrontal cortex—defining features of human cortical evolution. This also demonstrates that cortical neurogenesis, gliogenesis, and evolutionary expansion are not independent processes, but rather an integrated biological program coordinated by ERK, PKA, YAP/TAZ and SHH signaling.

During human corticogenesis, fRGs (full-span RGs) first undergo PyN genesis from GW8 to GW16. ^4,18,22^ By GW16, following approximately 9 weeks of neurogenesis, the human fetal cortex has undergone significant expansion and harbors a substantial population of PyNs that require glial cell-derived support and nutrient supply from the developing vascular network. Concurrently, YAP and SHH signaling, including CXCL12/CXCR signaling, progressively intensify in fRG cells. To accommodate the evolutionary pressure-driven expansion of PyN populations while concurrently meeting the heightened demand for glial cell production, human fRGs undergo a specialized division process, giving rise to oRGs and tRGs. ^4,18,22^ oRGs inherit the elongated basal processes from their fRGs and exhibit elevated levels of ERK and PKA activity, which are mutually reinforcing. ERK and PKA signaling enhancement suppresses gliogenic YAP and SHH signaling, thereby maintaining the neurogenic program in oRGs (Figure 7B). In contrast, tRGs inherit the apical membrane domains from fRGs, maintaining direct contact with the lateral ventricle. Subsequently, the enhanced activation of the YAP and SHH signaling orchestrates the establishment of E-tRGs and T-tRGs identities, steering their differentiation into ependymal cells and Tri-IPCs, respectively (Figure 7B, C).

The loss of FGFR or ERK function in RGs results in remarkably similar cortical phenotypes. ^17,66–68^ FGFR-mediated ERK signaling plays an essential role in multiple aspects of cortical development, including neural induction, cortical regional patterning, RG cell proliferation, self-renewal, survival, growth, and promoting *BMP7* and *HOPX* expression. ^15,17,30,32,33,66–68^ In this study, we observed that ERK and PKA signaling establish a mutually reinforcing positive feedback loop, mediated through repressing gliogenic YAP and SHH signaling, which collectively drive the expansion of the cortical RG population and extend the neurogenic period. Both the ERK and PKA signaling pathways are tightly regulated by robust negative feedback mechanisms, which effectively prevent uncontrolled signal amplification and maintain cellular signaling homeostasis. ^42,43^ Interestingly, this regulatory program of FGF-ERK signaling exhibits remarkable evolutionary conservation, with homologous mechanisms identified in hemichordates and amphioxus, suggesting its origin dates back at least 500 million years. ^78^ More importantly, FGF-ERK signaling exhibits a rostral-high to caudal-low gradient in the cortex, which promotes the expansion of the prefrontal cortex during evolution—a hallmark feature of the human cerebral cortex. Based on these findings, we propose that ERK signaling occupies a superior position in the regulatory hierarchy of signaling pathways governing cortical development and its surface-area expansion during evolution.

This study further led to the establishment of a novel model, in which a tripartite signaling network, centered on cross-repressive interactions among the ERK/PKA, YAP, and SHH pathways, regulates mammalian cortical neurogenesis, ependymal gliogenesis, and Tri-IPC genesis. Furthermore, it provides foundational evidence for the basic machinery underlying cortical expansion and evolution. Our model, however, inevitably challenges the conclusions of numerous previous studies, while our data support a distinct alternative perspective. For instance, although prior work has proposed that SHH signaling drives cortical expansion and gyrification, ^79–84^ our loss-of-function (*Smo-cko*) and gain-of-function (*dnPKA* overexpression) experiments demonstrate that SHH signaling instead plays a pivotal role in regulating the generation of cortical Tri-IPCs. Accordingly, we conclude that SHH signaling does not directly contribute to cortical expansion. Several previous studies have concluded that YAP signaling promotes cortical expansion and gyrification. ^85–89^ In contrast, our loss- and gain-of-function analyses demonstrate that YAP signaling primarily regulates cortical ependymal cell development via coordinated suppression of ERK, PKA, and SHH signaling pathways. Based on these findings, we conclude that YAP signaling does not drive cortical expansion. Numerous studies have proposed that human cortical RGs, particularly oRGs, generate cortical interneurons, OPCs, or Tri-IPCs. ^14,90–98^ In contrast, our findings argue against this view, as we propose that human oRGs predominantly produce PYNs. Cortical PyN fate specification is mediated by robust PKA and ERK signaling activity in RGs, which potently suppresses gliogenic YAP and SHH signaling pathways.

In summary, this study elucidates the molecular mechanisms responsible for the specification and lineage progression of distinct cortical RG subtypes across murine and human systems. It also uncovers fundamental principles that coordinate cortical neurogenesis, gliogenesis, and expansion, which are conserved during evolution. Thereby these findings provide critical insights into the fundamental mechanisms of cortical development (Figure 7).

## 4. Experimental Section

### Animals

All procedures involving animals were approved by and performed in accordance with the guidelines of the Fudan University Shanghai Medical College Animal Ethics Committee (No. 20230301-141). *Emx1-Cre* (JAX no. 005628), ^99^ *hGFAP-Cre* (JAX no. 004600), ^100^ *Rosa^Mek1DD^* (JAX no. 012352), ^101^ *Smo flox* (JAX no.004526), ^102^ *Map2k1* (exon 2) floxed and *Map2k2* (exon 4-9) floxed mice were described previously. ^17^ *SuperHippo* mice were provided by Professor Faxing Yu at Fudan University. ^65^ The day of detecting a vaginal plug was designated as E0.5. The day of birth was designated as P0. The sexes of the embryonic and early postnatal mice were not determined. For immunostaining analysis, normally, brains from 4-5 independent experiments were processed (n values refer to numbers of brains analyzed).

### Tissue preparation

Embryos were isolated from deeply anesthetized pregnant mice. Brains were dissected and fixed overnight in 4% paraformaldehyde (PFA) pre-treated with diethylpyrocarbonate (DEPC). Postnatal mice were deeply anesthetized and transcardially perfused with phosphate-buffered saline (PBS), followed by 4% PFA. All brains were post-fixed overnight in 4% PFA at 4°C and subsequently dehydrated in 30% sucrose for at least 24 hours. The brains were then embedded in O.C.T. compound (Sakura Finetek) and stored at −80°C. For analysis, mouse brains were sectioned into 20-μm-thick slices.

### Plasmid construction

*pCAG-ires-GFP* plasmid was from Addgene (Addgene #11150). The mouse *dnPKA* cDNA represents a dominant-negative form of PKA. ^103^ This variant is derived from PKA-RIa, which is encoded by the *Prkar1a* gene, and is characterized by mutations at the cAMP binding sites, rendering it unresponsive to cAMP. The *dnPKA* cDNA was subsequently cloned into the *pCAG-GFP* vector to generate *pCAG-dnPKA-ires-GFP* plasmids.

### *In Utero* Electroporation (IUE)

*In utero* electroporation was performed as described previously. ^13^ Briefly, pregnant mice were anesthetized using an animal anesthesia machine with isoflurane. A plasmid solution (final concentration: 1–2 μg/μl for each plasmid, 0.5 μl per embryo), mixed with 0.05% Fast Green (Sigma), was injected into the lateral ventricle of embryos using a beveled glass micropipette. Five electrical pulses (duration: 50 ms) were delivered at optimized voltages across the uterine wall, with a 950-ms interval between pulses, using a pair of 7-mm platinum electrodes (BTX, Tweezertrode 45-0488, Harvard Apparatus) connected to an electroporator (BTX, ECM830). The applied voltages were adjusted according to embryonic age: 33 V for E14, 35 V for E15, and 38 V for E18. Electroporated brains were analyzed at specified time points, as indicated in the main text.

### FlashTag labeling

FlashTag labeling was performed via in utero injection, and the FlashTag-labeled live cells were subsequently isolated using fluorescence-activated cell sorting (FACS). ^104,105^ To label embryonic day 15.5 (E15.5) cortical progenitors, FlashTag-CellTrace Yellow solution (Life Technologies, #C34567; 0.5 μl of 10 mM) was injected into the lateral ventricles of *SuperHippo* with or without *hGFAP-Cre* and *Emx1-Cre*, mice at E15.50, with analysis performed at E16.5. To label E17.0 progenitors, CellTrace Yellow solution was injected into the mouse lateral ventricles of *Smo^F/F^* and *hGFAP-Cre; Smo^F/F^* mice at embryonic day 17.0 (E17.0), with analysis performed at E18. To label P0 progenitors, the CellTrace solution was injected into the lateral ventricles of the following groups at postnatal day 0 (P0), with analysis conducted at P2: Control (*SuperHippo* without *Cre*), *hGFAP-Cre; SuperHippo*, *Smo^F/F^*, *hGFAP-Cre; Smo^F/F^*, *hGFAP-Cre; Map2k1/2-dcko*, and *Emx1-Cre; Map2k1/2-dcko* mice. Mice were sacrificed, and brains were immediately removed and submerged in fresh ice-cold Hanks’ balanced salt solution (HBSS; Gibco, 14175-095). Cortices were dissected, dissociated into single-cell suspensions, and subjected to FACS to purify FlashTag-labeled cells. Single-cell RNA sequencing (scRNA-Seq) was performed on FACS-sorted cells using the 10X Genomics platform.

### Immunohistochemistry

Immunohistochemistry was performed on 20-μm coronal sections of mouse brains using standard protocols. Sections were rinsed with Tris-buffered saline (TBS; 0.01 M Tris–HCl, 0.9% NaCl, pH 7.4) for 10 minutes, permeabilized with 0.5% Triton-X-100 in TBS for 30 minutes at room temperature (RT), and blocked in a solution of 5% donkey serum and 0.5% Triton-X-100 in TBS (pH 7.2) for 2 hours. After blocking, sections were incubated with primary antibodies diluted in blocking buffer overnight at 4°C. The following day, sections were washed three times with TBS (10 minutes each) and incubated with secondary antibodies (1:500; Jackson ImmunoResearch) for 2 hours in the dark at RT. After three additional TBS washes (10 minutes each), sections were counterstained with 4’,6-diamidino-2-phenylindole (DAPI; Sigma, 200 ng/mL) for 2 minutes. All primary antibodies utilized in this investigation, including their sources, catalog numbers, host species, and working dilutions, are detailed in Table S1.

### scRNA-Seq

Briefly, mouse cortices were dissociated into single-cell suspensions, and selected samples were screened by FACS prior to single-cell sequencing. scRNA-Seq libraries were generated using the Chromium droplet-based platform (10X Genomics) according to the manufacturer’s instructions (manual document part number: CG00052 Rev C). Purified cDNA libraries were quantified using an Agilent 2100 Bioanalyzer and sequenced on an Illumina NovaSeq 6000 platform. This study generated and systematically characterized a collection of 11 novel scRNA-Seq datasets, the complete details of which are presented in Table S2.

scRNA-Seq data derived from human cortical tissues at GW22, GW23, and GW26 were obtained from a previously published study. ^29^ For detailed methodological procedures, we refer readers to the original publication. ^29^

### scRNA-Seq data analysis

scRNA-Seq reads were aligned to the mm10 reference genome and quantified using ‘cellranger count’ (10x Genomics, v.7.1.0). Count data was further processed using the ‘Seurat’ R package (v.5.1.0). Genes detected in fewer than three cells, cells with fewer than 500 detected genes, and cells with more than 10% mitochondrial gene content were filtered out. The selection of highly variable genes was performed using the Variance Stabilizing Transformation (VST) method, which models mean-dependent technical noise and is robust to the influence of low-complexity cells. After filtering, the number of cells in each dataset was as follows:

E15.0 cortex: *Map2k1/2* (without *Cre*) 12,128 cells, 2,673 genes/cell, *Emx1-Cre; Map2k1/2-dcko* mice, 9,291 cells, 2,850 genes/cell; E16.5 FACS cortical progenitors: wild-type 14,800 cells, 1,504 genes/cell, *hGFAP-Cre; SuperHippo* 13,171 cells, 1,544 genes/cell; E16.5 FACS cortical progenitors: *SuperHippo* without *Emx1-Cre* 13,194 cells, 2,582 genes/cell, *Emx1-Cre; SuperHippo* mice, 14,641 cells, 2,616 genes/cell; E18.0 FACS cortical progenitors labeled by dnPKA-GFP in *hGFAP-Cre; Smo^F/F^* 11,508 cells, 2,369 genes/cell; P2 FACS cortical progenitors: control (*Smo^F/F^*), 14,292 cells, 2,498 genes/cell; *hGFAP-Cre; Smo^F/F^* mice, 7,694 cells, 2,681 genes/cell; control (*SuperHippo* heterozygotes without *hGFAP-Cre*), 12,054 cells, 2,258 genes/cell; *hGFAP-Cre; SuperHippo* mice, 11,539 cells, 2,120 genes/cell. These 11 novel scRNA-Seq data have been deposited in the GEO under the accession number GSE293205.

Gene expression data were normalized using the global-scaling method “LogNormalize”. The top 2,000 highly variable genes were identified with the FindVariableFeatures function using the vst method. To minimize the influence of the cell cycle on clustering and dimensionality reduction, cell cycle-associated gene sets were used to score the cell phase of each cell with the CellCycleScoring function, and the difference between G2M and S phase scores was regressed out using the ScaleData function. The scaled z-scored residuals were used for Principal Component Analysis (PCA). Statistically significant principal components, identified through a resampling test, were retained for FindNeighbors and FindClusters analysis. Uniform Manifold Approximation and Projection (UMAP) was used for visualization of cell clustering. Differentially expressed genes (DEGs) among clusters were identified using the Wilcoxon rank sum test, comparing cells in each cluster against all other cells.

### Image acquisition and analysis

All brain section images in this study were acquired using an Olympus VS120 Automated Slide Scanner (10X, 20X) and an FV3000 confocal microscope system (40X). Images were processed using Adobe Photoshop for clarity, false colorization, and overlay as needed. Both Adobe Photoshop and Adobe Illustrator were used to adjust images without altering the original data.

### Quantification and statistical analysis

Quantitative images were acquired using the Olympus VS120 Automated Slide Scanner (10X or 20X). Statistical analyses were performed using GraphPad Prism 10 and IBM SPSS Statistics 29.0. For each experiment, at least four control or mutant mouse samples were analyzed. The following quantifications were performed:

1. Quantification of GSX2-, OLIG2-, and EGFR-positive cells in the cortical VZ/SVZ (width: 1200 pixels; height: 800 pixels) at E18 in *Smo^F/F^*, *Smo^F/F^+ dnPKA*, *hGFAP-Cre; Smo^F/F^*, and *hGFAP-Cre; Smo^F/F^+ dnPKA* mouse brains (see Figure 4D, E).
2. Quantification of FOXJ1- and CRYAB-positive cells in the cortex of control and *Emx1-Cre; Map2k1/2-dcko* mice at P0. FOXJ1- and CRYAB-positive cells in the dorsal cortical VZ/SVZ were counted. Four brain slices from corresponding locations were analyzed per sample (see Figure 5D, E).
3. Quantification of GSX2^+^ and OLIG2^+^ cells in the cortex of control and *hGFAP-Cre; SuperHippo* mice. Four brain slices from corresponding positions were selected per sample, and GSX2^+^ and OLIG2^+^ cells in the entire VZ/SVZ region were counted (see Figure 3D, E, and Figure S5D, E, Supporting Information).
4. Quantification of cortical area at P21 in control and *Emx1-Cre; Map2k1/2-dcko* mice. Six brain slices from corresponding positions were analyzed per sample, and the cortical area was measured using Photoshop (see Figure S9A-C, Supporting Information).
5. Quantification of EOMES^+^ cells in the cortical VZ (width: 300 pixels; height: 300 pixels) of control and *Emx1-Cre; Rosa^Mek1DD^* mice at E13.5. Four brain slices from corresponding positions were analyzed per sample (see Figure S9D, E, Supporting Information).
6. Quantification of cortical area at P76 in control and *Emx1-Cre; Rosa^Mek1DD^* mice. Six brain slices from corresponding positions were analyzed per sample, and the cortical area was measured using Photoshop (see Figure S9F-H, Supporting Information).
7. Quantification of FOXJ1- and CRYAB-positive cells in the cortex of control and *Emx1-Cre; Map2k1/2-dcko* mice at P2. FOXJ1- and CRYAB-positive cells in the dorsal cortical VZ/SVZ were counted. Four brain slices from corresponding locations were analyzed per sample (see Figure 6C, D). Data are presented as mean ± SEM. Statistical significance for single comparisons was determined using unpaired t-tests. A *P*-value < 0.05 was considered statistically significant (\**P* < 0.05, \*\**P* < 0.01, \*\*\**P* < 0.001).

## Additional information

Supporting Information is available from the Wiley Online Library or from the author.

## Acknowledgments

We sincerely thank Professor John L. Rubenstein for his insightful comments and critical editing of the manuscript. We are also grateful to Professors Arnold Kriegstein and Bin Chen for their valuable feedback and constructive critiques on our data. We thank Professor Faxing Yu for sharing the *SuperHippo* mice. This study was supported by the Ministry of Science and Technology of China (STI2030-2021ZD0202300), National Natural Science Foundation of China (NSFC 31820103006, 32070971, 32100768, 32200776, and 32200792), China Postdoctoral Science Foundation (2024T170179), Shanghai Municipal Science and Technology Major Project (No.2018SHZDZX01), ZJ Lab, and Shanghai Center for Brain Science and Brain-Inspired Technology.

## Author contributions

Conceptualization: Z.Y. Data curation: T.F., Z.Z., Z.X. and J.L. Funding acquisition: Z.Z., Z.X., T.M., and Z.Y. Investigation: Z.Z., Z.X., T.F., J.L., F.Y., C.Y., W.Z., Z.S., Y.G., M.S., Z.L., J.D., and X.L. Resources: Z.Y. Supervision: Z.Y. Writing: Z.Y. All of the authors contributed to reviewing and editing of the manuscript.

## Data Availability Statement

The datasets (scRNA-Seq from 11 samples, Table S2) generated during the current study are available in the Gene Expression Omnibus (GEO: GSE293205). The E18.0 *Smo^F/F^*(control) and *hGFAP-Cre; Smo^F/F^* mouse cortex scRNA-Seq data were used from our previous study (GSE221389). ^25^ The P2 *Emx1-Cre; Map2k1/2-dcko* and P2 *hGFAP-Cre; Map2k1/2-dcko* mouse cortex scRNA-Seq data were used from our previous studies (GSE274547). Bulk RNA-Seq data from the SVZ of wild-type control mice (n=3) and tumor cells derived from ∼P20 mice transfected with YAP-M at E14.5 (n=4), and ChIP-Seq and Cut&Run data for YAP1 and H3K27ac in both YAP-M-induced tumor cells and YAP1-expressing mouse neural stem cells were used in previous studies (GSE181867). ^61^ scRNA-Seq data derived from human cortical tissues at GW22, GW23, and GW26 were obtained from a previously published study (GSE162170). ^29^

## Supporting Information

**Figure S1.**
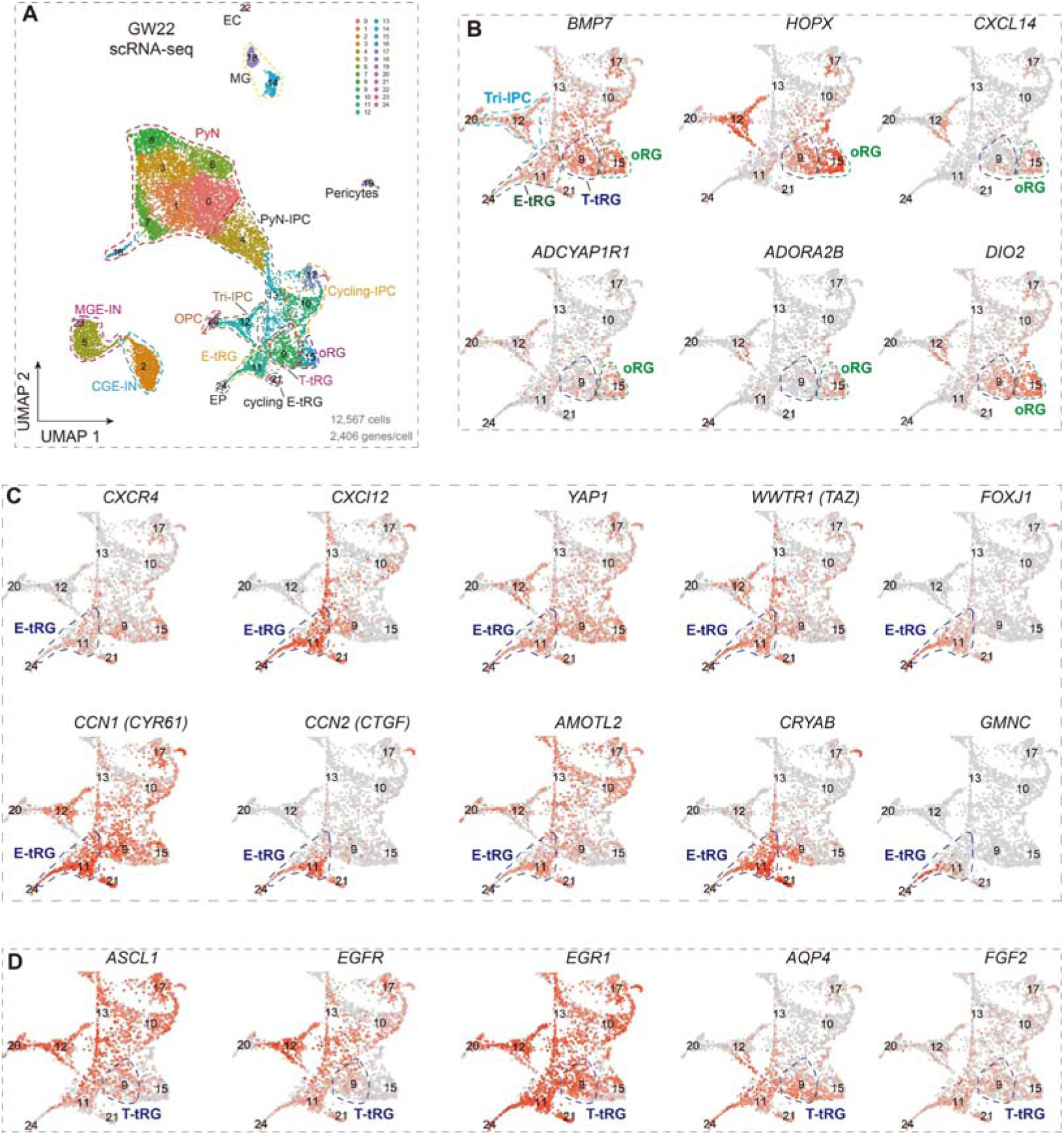
Expression of cell-type-specific markers in scRNA-Seq data at GW22. **A)** UMAP of GW22 scRNA-Seq cells colored by cluster (from Figure 1A). **B-D)** UMAP plots showing gene expression in human cortical progenitors. Note that key components of the cortical PKA signaling, including *ADCYAP1R1, ADORA2B, DIO2*, and *CXCL14*, demonstrate significantly elevated expression levels in oRGs (**B**). *FOXJ1* and *GMNC* (*GEMC1*) are two master regulators essential for initiating the multiciliation program. Note that the onset of FOXJ1 expression in E-tRGs precedes that of GMNC (**C**). oRG, outer radial glia; E-tRG, ependymocyte-generating truncated radial glia, T-tRG, Tri-IPC-generating tRG; EP, ependymal cell; Tri-IPCs, tripotential intermediate progenitor cells; OPC, oligodendrocyte-IPCs; PyN, cortical glutamatergic pyramidal neuron; PyN-IPC, PyN intermediate progenitor cells; CGE-IN, caudal ganglionic eminence interneuron; MGE-IN, medial ganglionic eminence interneuron; EC, endothelial cell; MG, microglia.

**Figure S2.**
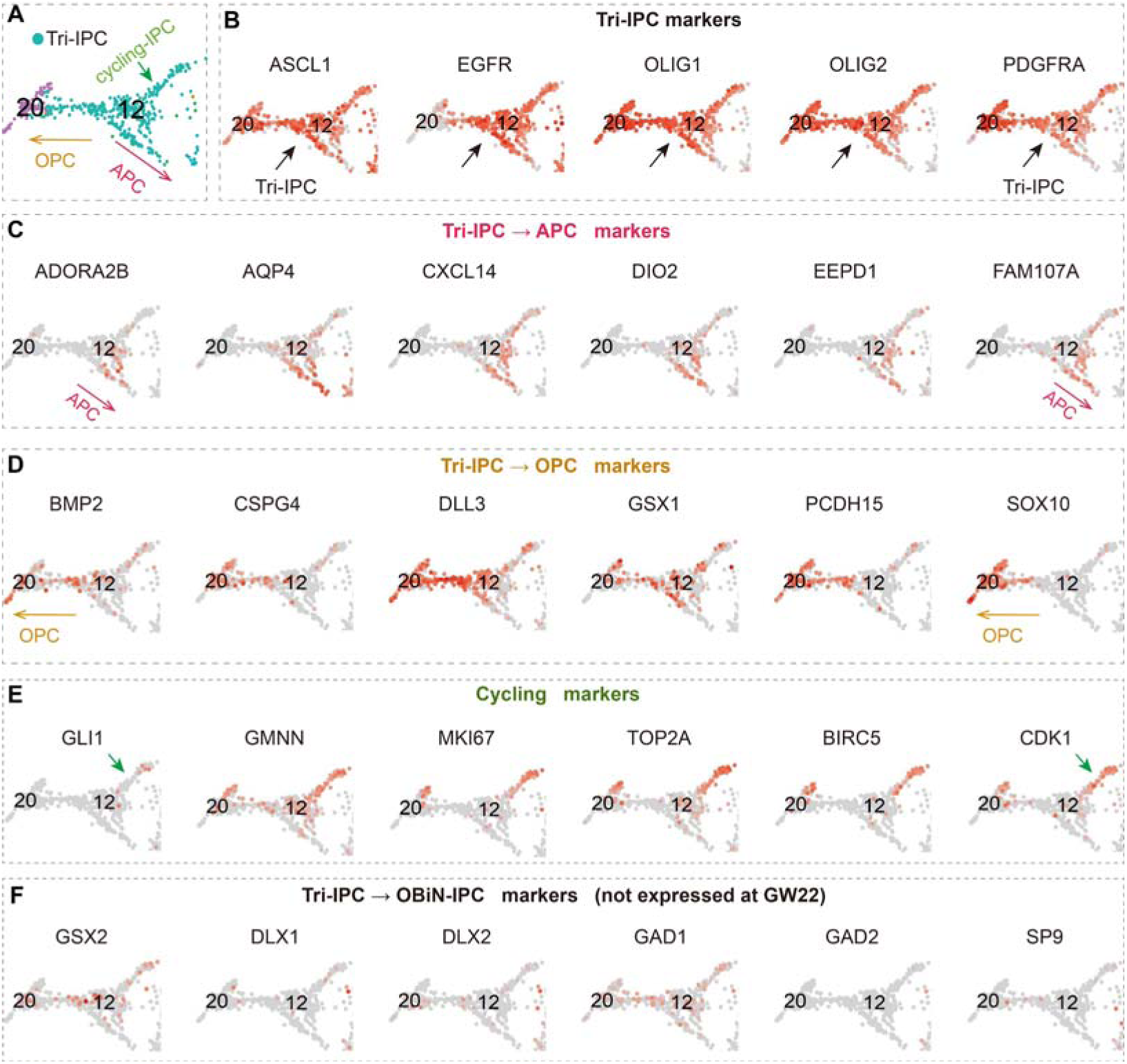
Cortical T-tRG-derived Tri-IPCs at GW22 do not generate OBIN-IPCs. **A-F)** While marker genes for cortical Tri-IPCs, APCs, OPCs, and cycling progenitors, are identified (arrows in **A**-**E**), OBIN-IPC marker genes are not observed in Tri-IPC clusters at GW22 **F)**. APC, astrocyte-IPCs; OBIN-IPC, IPCs for cortically derived olfactory bulb interneuron.

**Figure S3.**
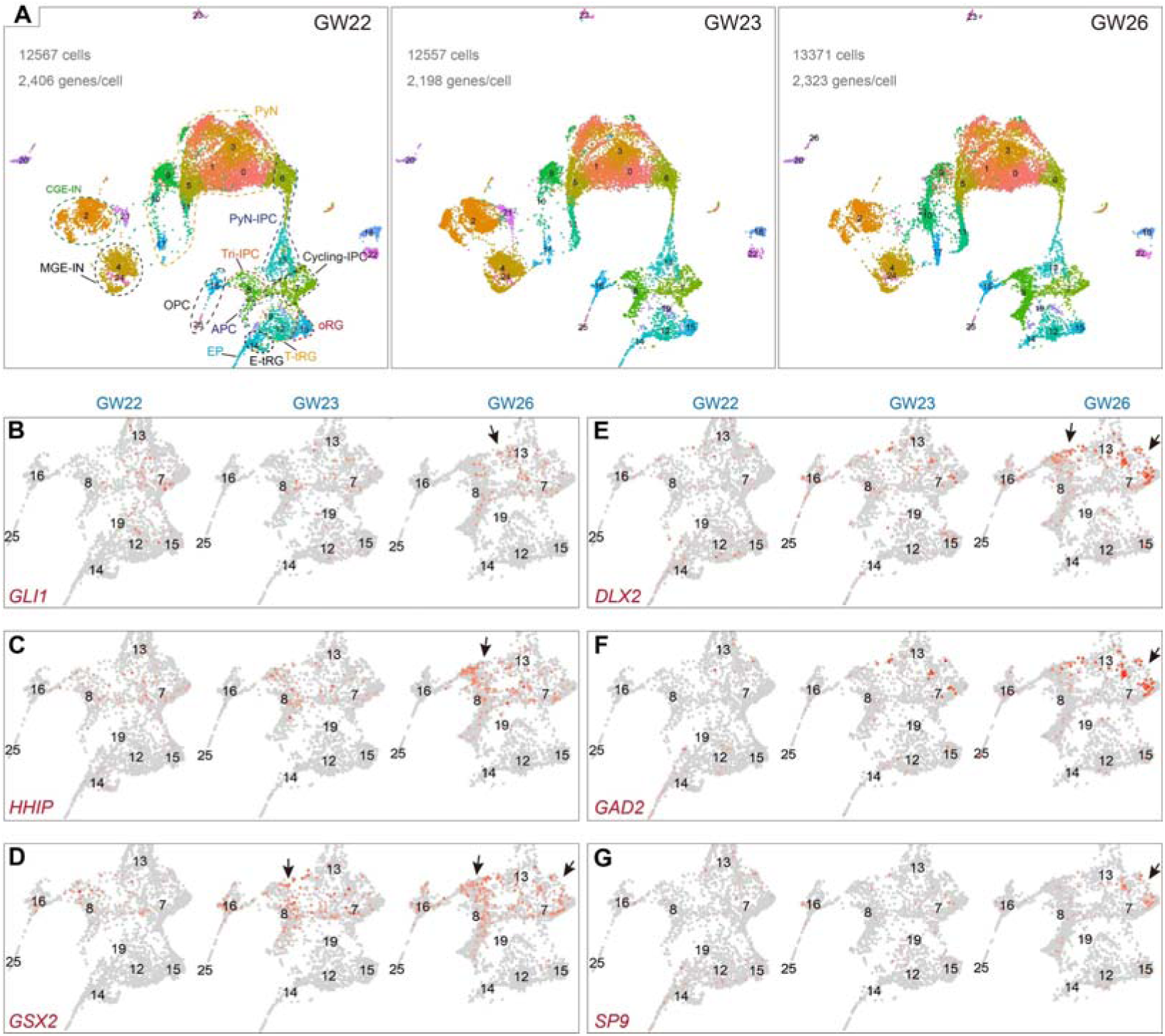
Cortical T-tRG-derived Tri-IPCs at GW23 and GW26 undergo progressive generating OBIN-IPCs. **A)** UMAP of GW22, GW23, and GW26 scRNA-Seq cells colored by cluster. **B-C)** Expression levels of SHH signaling markers *GLI1* and *HHIP* (arrows) gradually increase in cortical progenitors from GW22 to GW26. **D-G)** Expression of OBIN-IPC markers, including *GSX2, DLX2, GAD2*, and *SP9* within Tri-IPC cluster (arrows), are observed at GW26.

**Figure S4.**
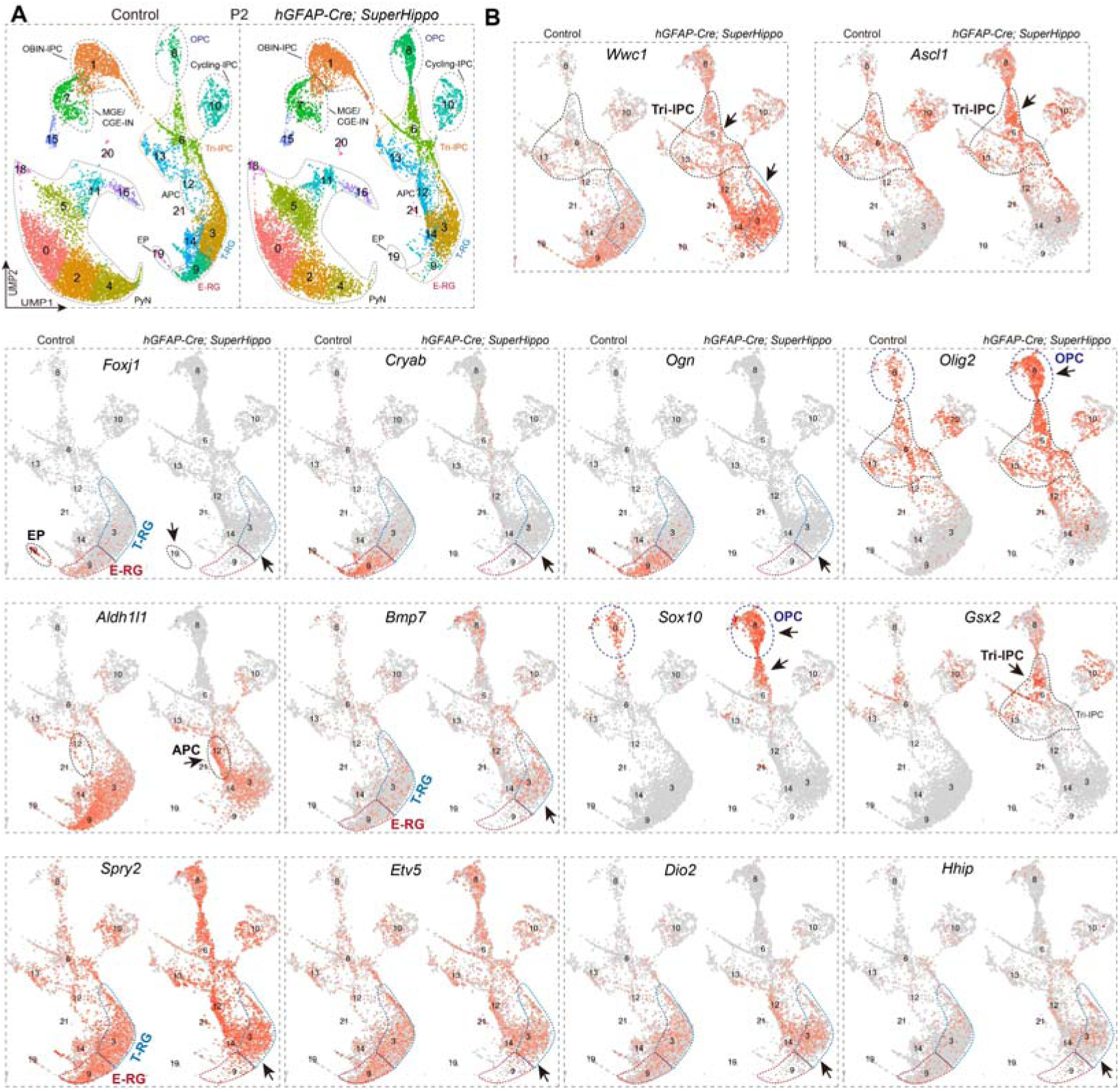
Loss of YAP signaling results in the depletion of cortical E-RGs and ependymal cells. **A)** UMAP of scRNA cells colored by cluster (from Figure 3A). **B)** UMAP plots showing *Wwc1* gene expression was increased in *hGFAP-Cre; SuperHippo* (*Wwc1*-derived *SuperHippo* minigene) mice at P2 (arrows). Note that loss of YAP signaling results in the depletion of cortical E-RGs and the loss of early ependymal marker gene expression in cortical T-RGs, including *Foxj1, Cryab*, and *Ogn* (arrows). In contrast, there is the upregulation of genes related to Tri-IPCs (*Olig2*), APCs (*Aldh1l1*), OPCs (*Sox10*), OBIN-IPCs (*Gsx2*), ERK signaling (*Bmp7*, *Spry2*, and *Etv5*), PKA signaling (*Dio2*), and SHH signaling (*Hhip*).

**Figure S5.**
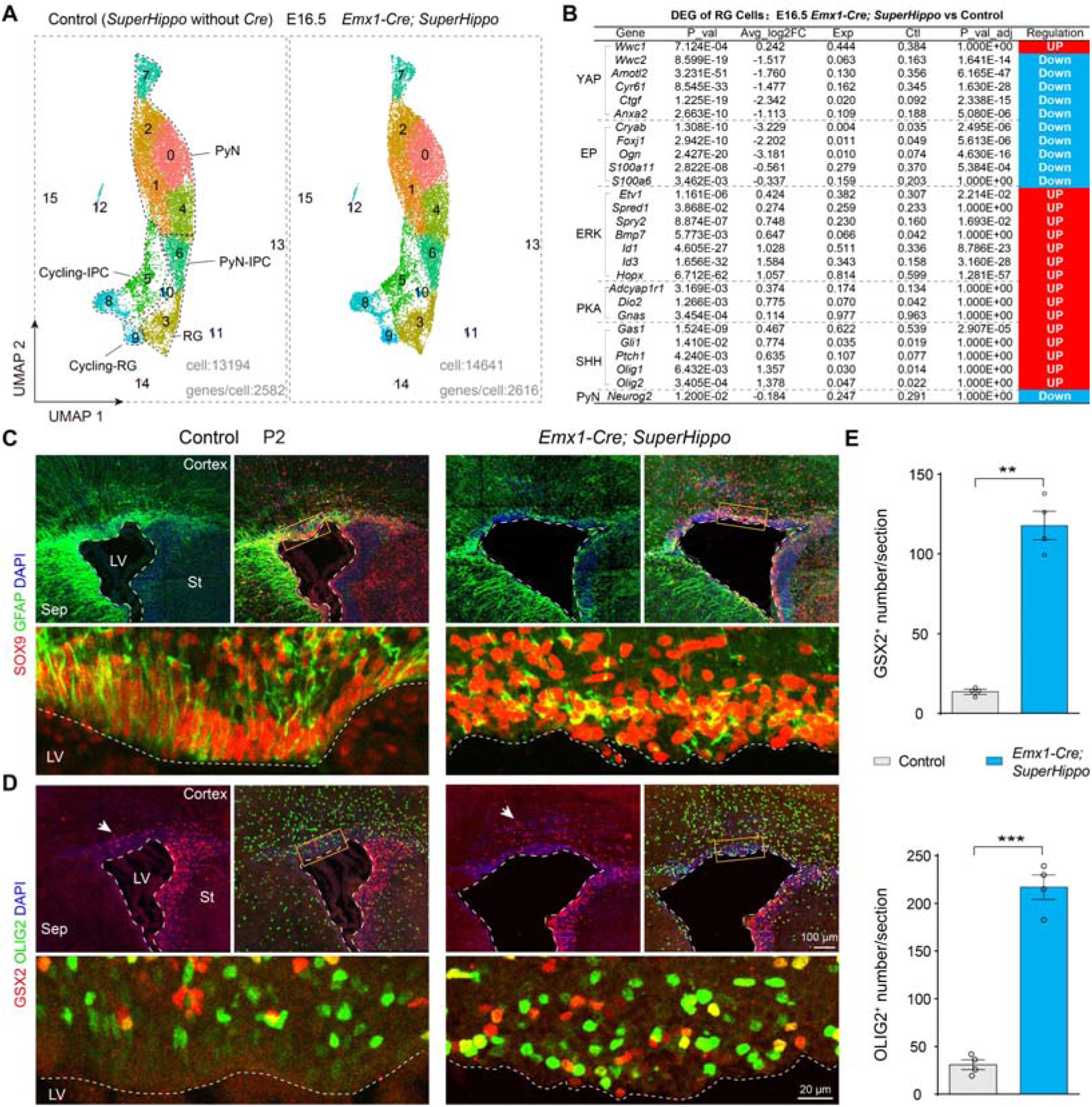
YAP signaling represses ERK, PKA, and SHH pathway activities in cortical RGs. **A)** UMAP of scRNA cells colored by cluster. **B)** scRNA-Seq analysis revealed differentially expressed genes (DEG) in E16.5 cortical RGs (cluster 3 in **A**) of *Emx1-Cre; SuperHippo* mice relatively to controls (*SuperHippo* mice without *Emx1-Cre*). **C-E)** At P2, immunostaining reveals disruption of the cortical ventricular surface and the loss of apical SOX9- and GFAP-expressing immature ependymal cells in the *Emx1-Cre; SuperHippo* cortex. Consistent with the elevated SHH signaling activity in cortical RGs, quantitative analysis demonstrated a significant expansion of GSX2- and OLIG2-expressing progenitor in the cortical VZ/SVZ of E*mx1-Cre; SuperHippo* mice.

**Figure S6.**
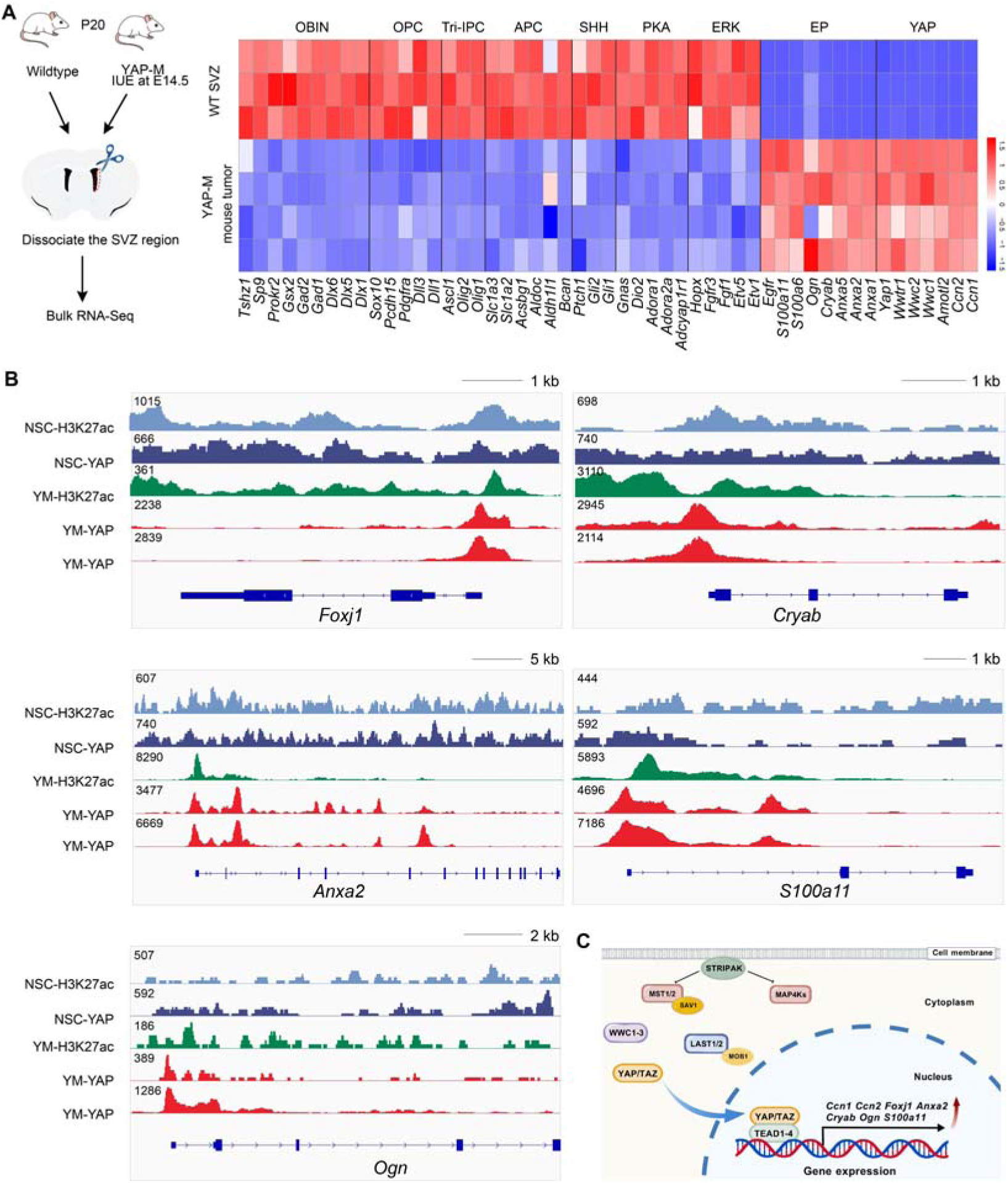
YAP signaling promotes the cortical E-RG cell lineage by directly activating key transcriptional regulators of ependymal development. **A)** Bulk RNA-Seq reanalysis. Heatmap of differentially expressed genes between the SVZ cells of wild-type control mice (n=3) and the tumor cells derived from ∼P20 mice transfected with YAP-M at E14.5 (n=4 independent replicates). Note that the YAP-M fusion oncogene markedly upregulated YAP signaling components and early ependymal markers, while strongly suppressing key pathway genes (ERK, PKA, SHH) and cortical progenitor markers (Tri-IPCs, APCs, OPCs, OBIN-IPCs). **B, C)** ChIP-Seq analysis showing YAP1 binding and H3K27ac modification profiles on early ependymal cell marker gene loci in YAP-M tissue and YAP-expressing wild type neural stem cells from P20 mice.

**Figure S7.**
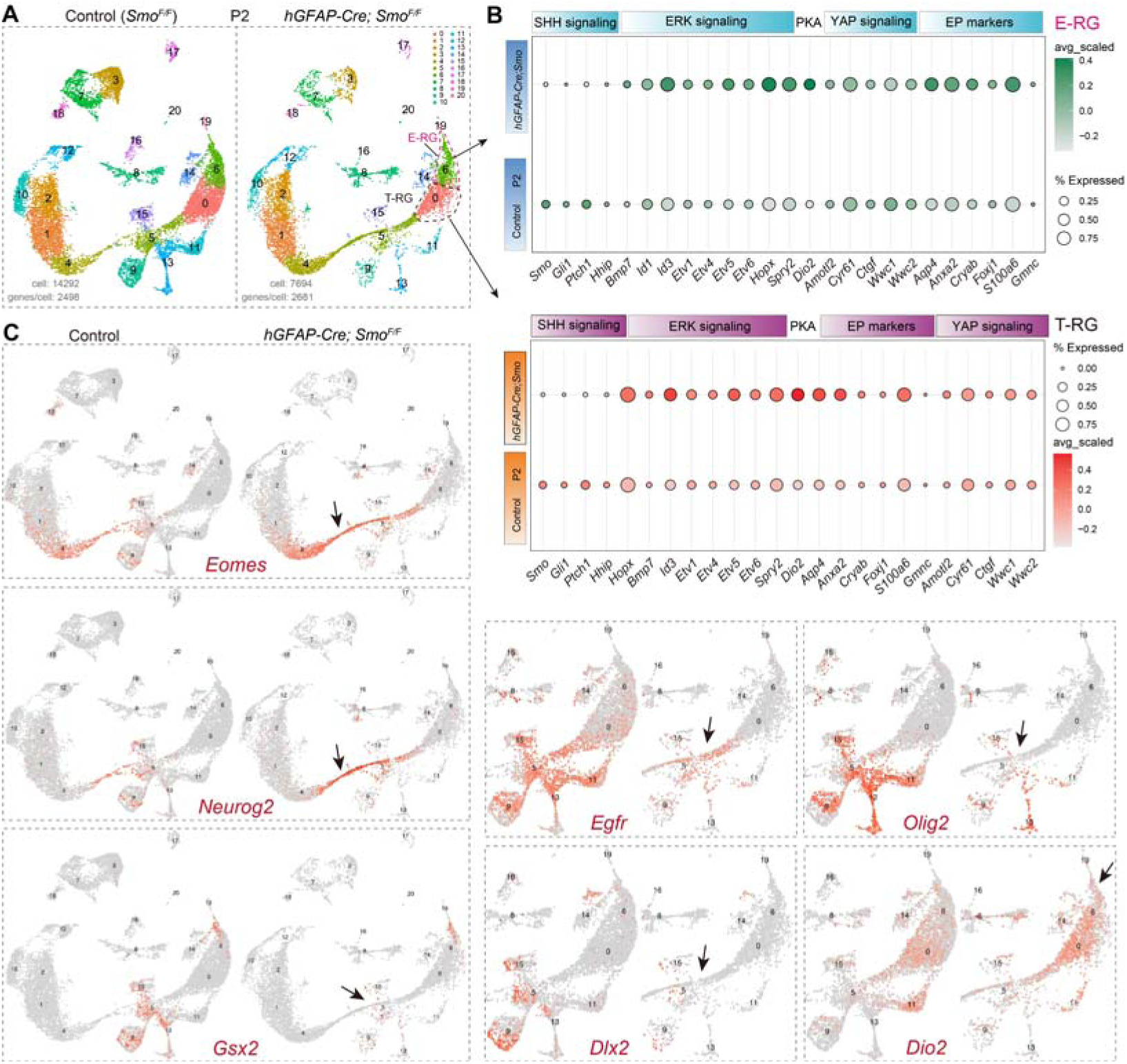
SHH-SMO signaling inhibits PKA, ERK, and YAP signaling pathways. **A)** UMAP of scRNA cells colored by cluster. **B)** Bubble plot showing differentially expressed genes (DEG) in cortical E-RGs and T-RGs in control (*Smo^F/F^*) and *hGFAP-Cre; Smo^F/F^* mice at P2. **C)** UMAP plots showing differentially expressed genes in the cortex of control (*Smo^F/F^*) and *hGFAP-Cre; Smo^F/F^* mice at P2. In the absence of *Smo* function, *Eomes, Neurog2* and *Dio2* expression were increased in cortical RGs and/or progenitors, whereas *Gsx2*, *Egfr*, *Olig2*, and *Dlx2* expression (arrows) were significantly reduced.

**Figure S8.**
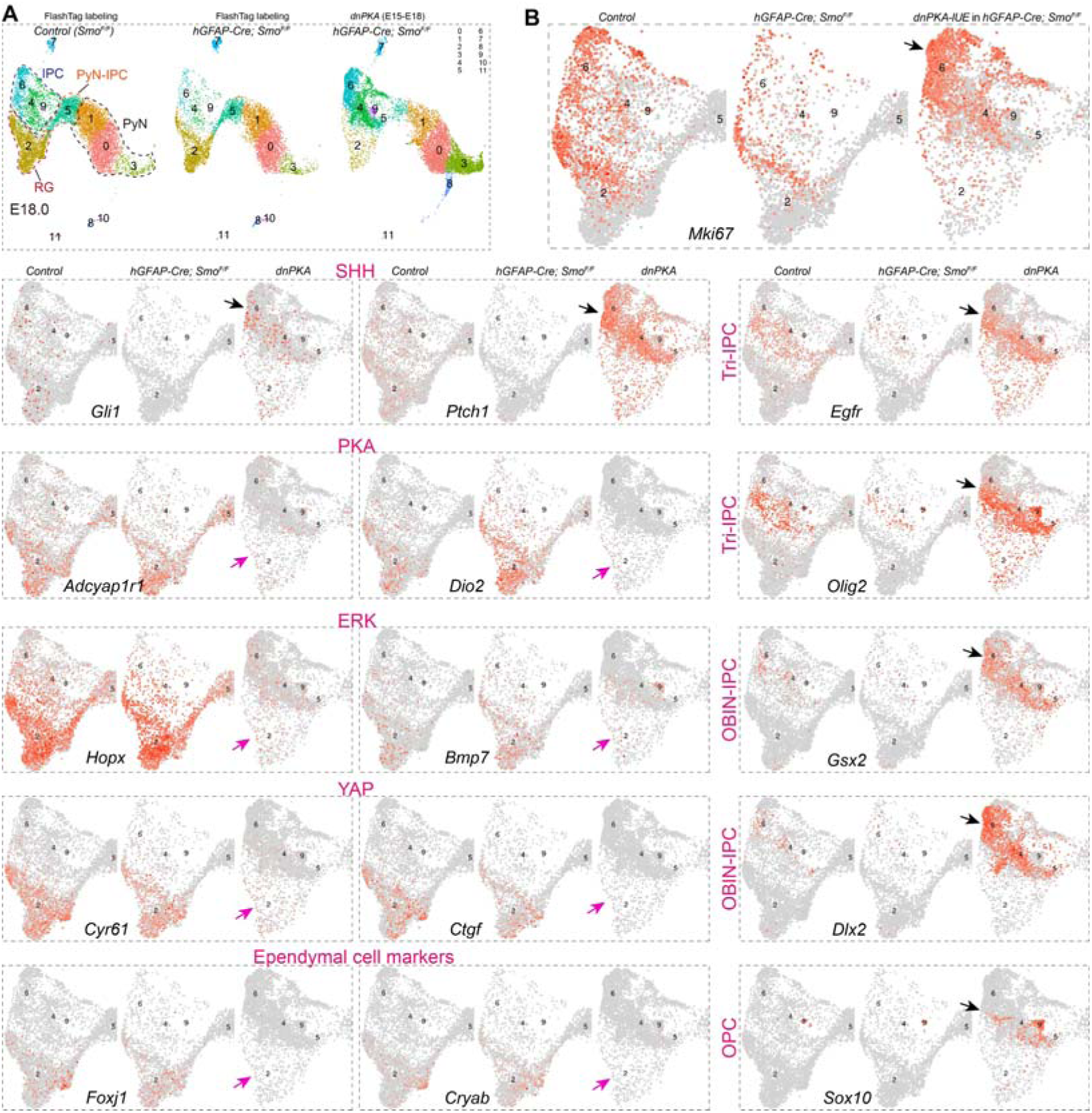
PKA signaling strongly inhibits SHH-SMO signaling. **A)** UMAP of scRNA cells colored by cluster (from Figure 4B). **B)** UMAP plots showing individual gene expression in the cortex of *Smo^F/F^, hGFAP-Cre; Smo^F/F^,* and *hGFAP-Cre; Smo^F/F^* mouse at E18.0 with IUE of *dnPKA* at E15.0. The complete loss of PKA in cortical RGs resulted in a significant increase in SHH signaling (*Gli1* and *Ptch1*), even in the absence of *Smo* function. This is accompanied by the depletion of PKA (*Adcyal1r1* and *Dio2*), ERK (*Hopx* and *Bmp7*), and YAP (*Cyr61* and *Ctgf*) signaling, as well as the loss of early ependymal marker gene (*Foxj1* and *Cryab*) expression. In contrast, there is a significant upregulation of genes associated with Tri-IPCs (*Egfr* and *Olig2*), OBIN-IPCs (*Gsx2* and *Dlx2*), and OPCs (*Sox10*) (arrows).

**Figure S9.**
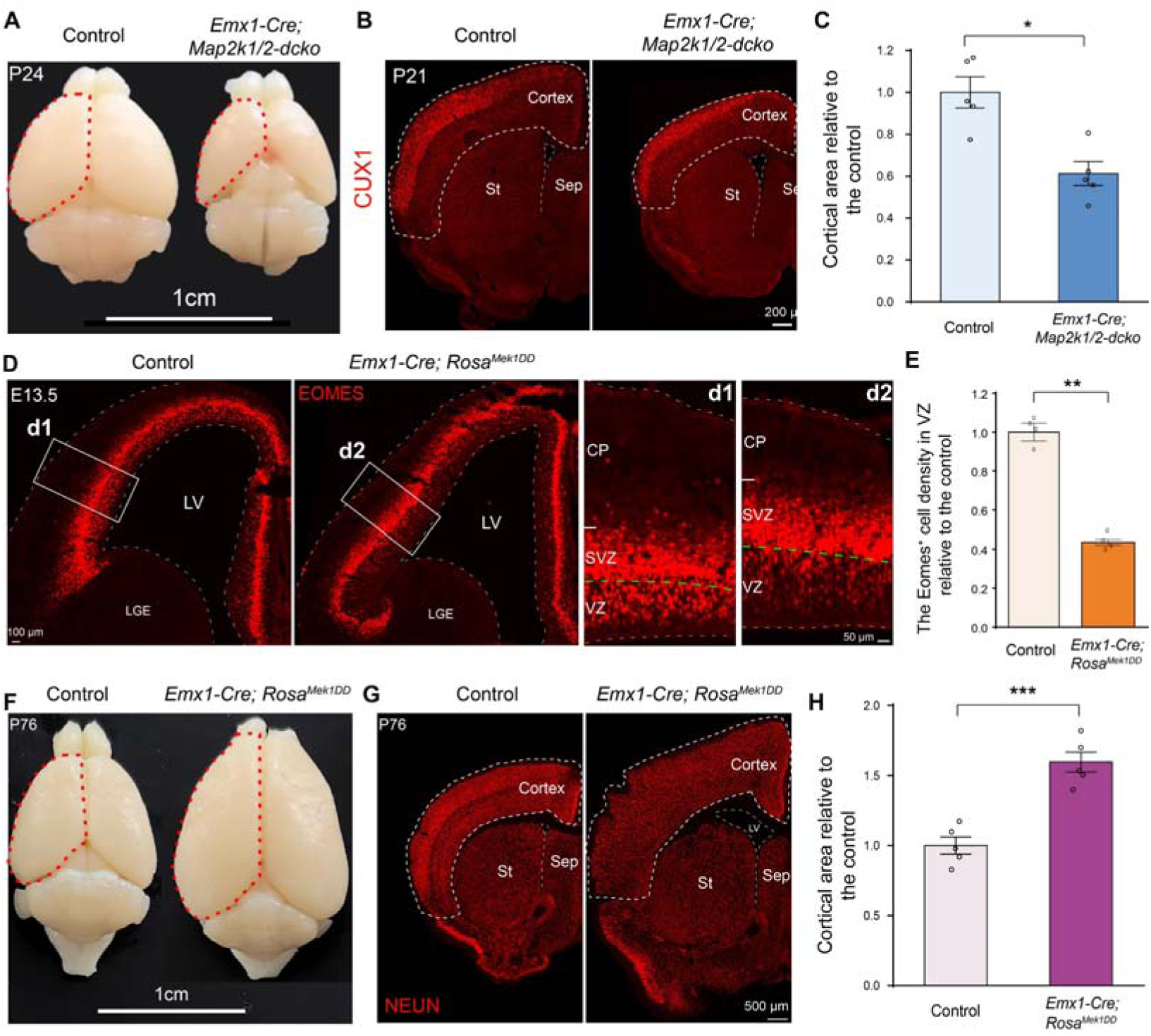
Loss and gain of ERK function from E10.5 cortical RGs result in pronounced microcephaly and macrocephaly, respectively. **A)** Whole mount images of control and *Emx1-Cre; Map2k1/2-dcko* brains. **B)** Immunostaining for CUX1 revealed fewer upper-layer PyNs and a smaller cortical area in *Emx1-Cre; Map2k1/2-dcko* cortices. **C)** Quantification of the cortical area within the cortical plate in control and *Emx1-Cre; Map2k1/2-dcko* mice. **D,E)** Immunostaining for EOMES, a marker of PyN-IPCs, revealed fewer EOMES-expressing cells in the VZ of *Emx1-Cre; RosaMEK1DD* mice at E13.5, suggesting an expansion of cortical RGs and restricted neuronal differentiation following enhanced ERK signaling. LGE, Lateral ganglionic eminence. **F)** Whole mount images of control and *Emx1-Cre; RosaMEK1DD* brains at P76. **G)** Immunostaining for NEUN, a marker of neurons. **H)** Quantification revealed a larger cortical area in *Emx1-Cre; RosaMEK1DD* mice at P76.

**Figure S10.**
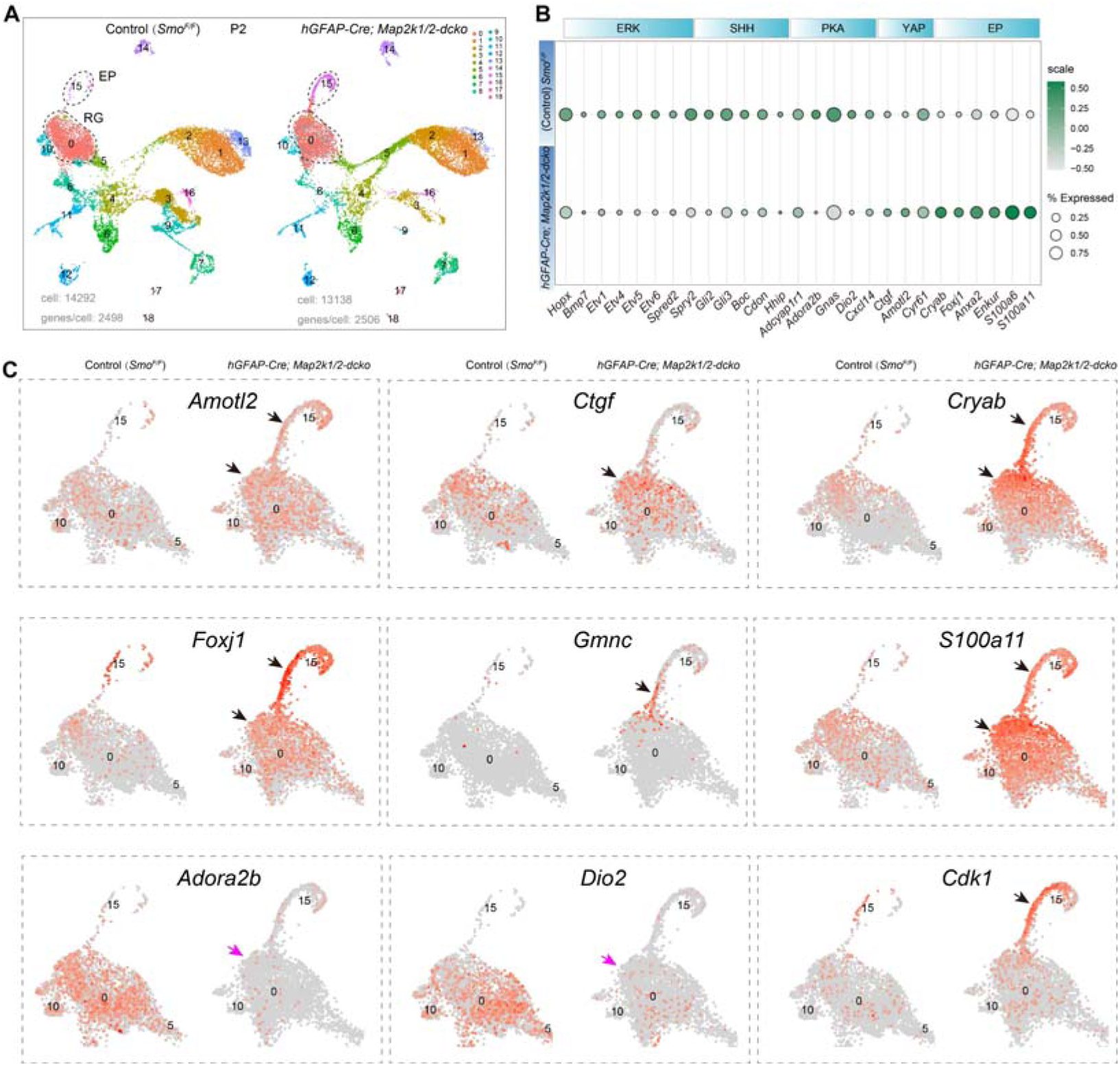
ERK signaling sustains PKA signaling and inhibits YAP signaling in cortical RGs. **A)** UMAP showing annotated cell clusters based on scRNA-Seq analysis of P2 control and *hGFAP-Ce; Map2k1/2-dcko* cortical cells. **B)** Bubble plot showing differentially expressed genes in cortical RGs (cluster 0 in **A**) of *hGFAP-Cre; Map2k1/2-dcko* mice compared to controls at P2. **C)** UMAP plots showing differentially expressed genes of YAP and PKA signaling pathway and early ependymal cell marker genes.

**Table S1.** Antibodies used in this study.

| primary Ab | Species | Dilution | Manufacturer | Cat. No. |
| --- | --- | --- | --- | --- |
| CUX1 | Rabbit | 1:200 | Santa Cruz | sc-13024 |
| CRYAB | Mouse | 1:500 | Abcam | ab13496 |
| EGFR | Goat | 1:1000 | R&D System | BAF1280 |
| EOMES | Guinea pig | 1:500 | Oasis Biofarm | OB-PGP022 |
| FOXJ1 | Mouse | 1:1000 | Invitrogen | 14-9965-80 |
| GFP | Chicken | 1:3000 | Aves labs | GFP-1020 |
| GFAP | Rabbit | 1:1000 | Dako | Z0334 |
| GSX2 | Rabbit | 1:500 | Millipore | ABN162 |
| NEUN | Rabbit | 1:1000 | Biosensis | R-3770-100 |
| OLIG2 | Rabbit | 1:500 | Millipore | AB9610 |
| OLIG2 | Rat | 1:500 | Oasis Biofarm | OB-PRB009 |
| pERK1/2 | Rabbit | 1:400 | Cell Signaling | #4370 |
| SOX9 | Rabbit | 1:1000 | Abcam | ab185966 |
| YAP/TAZ | Rabbit | 1:1500 | Cell Signaling | 14074S |

**Table S2.** Single-cell RNA sequencing (scRNA-Seq) was performed on 11 mouse cortical samples newly generated for this study.

| Number | Age | Genotype | Notes |
| --- | --- | --- | --- |
| 1 | E15.0 | *Map2k1/2* (without *Cre*) | Whole cortex |
| 2 | E15.0 | *Emx1-Cre; Map2k1/2-dcko* | Whole cortex |
| 3 | E16.5 | *Wild-type littermate* | FlashTag labeling at E15.5 |
| 4 | E16.5 | *hGFAP-Cre; SuperHippo* | FlashTag labeling at E15.5 |
| 5 | E16.5 | *SuperHippo* (without *Cre*) | FlashTag labeling at E15.5 |
| 6 | E16.5 | *Emx1-Cre; SuperHippo* | FlashTag labeling at E15.5 |
| 7 | E18.0 | *hGFAP-Cre; Smo<sup>F/F</sup> + dnPKA* | IUE (*dnPKA*) labeling at E15.0 |
| 8 | P2 | *SuperHippo* (without *Cre*) | FlashTag labeling at P0 |
| 9 | P2 | *hGFAP-Cre; SuperHippo* | FlashTag labeling at P0 |
| 10 | P2 | *Smo<sup>F/F</sup>* | FlashTag labeling at P0 |
| 11 | P2 | *hGFAP-Cre; Smo<sup>F/F</sup>* | FlashTag labeling at P0 |

